## Supplementary material for "Tuning apicobasal polarity and junctional recycling in the hemogenic endothelium orchestrates the morphodynamic complexity of emerging pre-hematopoietic stem cells": Torcq et al SUPPLEMENTAL TEXT + FIGURES

#### Legends to supplement figures

**Figure 1 – figure supplement 1. A**, 2 EHT pol+ cells visualized using a *Tg(Kdrl:Gal4;UAS:RFP;4xNR:eGFP-podxl2)* embryo and time-lapse sequence (initiated at 55 hpf) obtained with spinning disk confocal microscopy. The images (single z-planes) are extracted from the same time lapse sequence than the one used for ***Figure 1C***. Green channel (eGFP-podxl2) only is shown. Green arrows point at the evolution of the connection between the aortic/eht cell lumens at t = 0, 10, 30 min. Asterisks label the lumen delimited by the luminal/apical membranes. Bar = 8$\mu$m. **B**, model summarizing and interpreting the temporal evolution of the luminal/apical membrane (in green, the asterisks mark the lumen of the apparent vacuole-like intracellular membrane structures) after the release of an EHT pol+ cell from the aortic floor. **Step 1**, the pseudo-vacuole filled with fluid and delimited by eGFP-podxl2 as visualized at t = 65 min in **A** (and also at 45 min in ***Figure 1C***, 2 asterisks) is consumed partly via budding (after sorting of eGFP-podxl2); these budding profiles can be seen on the left image (pink arrowheads) corresponding, for eht cell 1, to the plane 16 of the time point t = 80 min of the time lapse sequence (see the Z-stack in ***Figure 1 – video 2***). **Step 2**, after sorting and budding, the cell remains with pseudo-endocytic Podxl2 containing membranes and the remaining vacuolar structures filled up with fluid regress, putatively by chasing water as illustrated in **step 3**. **Step 4**, the eht cell remains with pseudo-endocytic Podxl2 containing membranes that label newly born precursors of HSPCs. Note that since the cell remains in contact with the aortic floor while the pseudo-vacuole is regressing, the vacuole-like intracellular membrane proximal to aortic cells may never undergo fission (see ***Figure 1A***, steps 4-4') but gets consumed via budding and flattening upon water chase (steps 1-3 that are similar to ***Figure 1A***, steps 3-3').

**Figure 1 – figure supplement 2. EHT pol+ and EHT pol- cells recover their respective morphology after mitosis. A**, EHT pol+ cell extrusion and division visualized using a *Tg(Kdrl:Jam3b-eGFP;Kdrl:nls-mKate2)* embryo and time-lapse sequence (initiated at 55 hpf) obtained using laser scanning confocal microscopy (single z-planes). This transgenic line allows the visualization of nuclei (nls-mKate2, magenta) and intercellular contacts as well as plasma membranes (eGFP-Jam3b, see ***Methods*** and ***Figure 5*** for more information). The eht 1 cell (white box) starts invaginating before t = 0 min, and displays the characteristic luminal membrane invagination of EHT pol+ cells at t = 40 min. At t = 60 min, the cell undergoes mitosis. At t = 80min, the two daughter cells (eht 1' and eht 1'') are still contacting each other. At this point, they still have the slightly rounded morphology of post-mitotic cells. At t = 200 min, the eht 1' cell has resumed its extrusion and its luminal membrane is invaginated. By t = 340 min, the extrusion of the eht 1' cell is complete, and the cell remains in contact with the aortic floor. The eht 1'' cell resumes its extrusion later, also displaying an invagination of its luminal membrane (but not visible on the same plane as for the eht 1' cell). White asterisks label the lumen delimited by the luminal/apical membranes. Bar = 10$\mu$m. **B**, EHT pol- cell extrusion and division visualized using a *Tg(Kdrl:Gal4;UAS:RFP;4XNR:eGFP-podxl2)* embryo and time-lapse sequence (initiated at 54 hpf) obtained with spinning disk confocal microscopy (single z-planes), at the bottom of the trunk region. The eht 1 cell (white box) is undergoing extrusion at t = 0 min, and displays the characteristic rounded/ovoid shape of EHT pol- cells at t = 0 min and t = 10 min. At t = 20 min, the cell starts undergoing mitosis. At t = 130 min, we can observe the two daughter cells (eht 1' and eht 1'') still contacting each other. From t = 130 min to t = 235 min, the two daughter cells slowly continue their extrusion, keep contacting each other, and increase their roundness, with partitioning of eGFP-podxl2 between the luminal and basal membranes. All throughout the time-lapse sequence, the eht 1 cell and, subsequently, the eht 1' and eht 1'' cells do not display any enrichment of eGFP-podxl2 at their luminal membranes.

**Figure 1 – figure supplement 3. EHT pol+ and EHT pol- cells express eGFP driven by the CD41 promotor.** Single z-plane of a 55 hpf *Tg(Kdrl:mKate2-podxl2;CD41:eGFP)* embryo imaged using spinning disk confocal microscopy, in the trunk region. Top panel, left box (with 2X magnification beneath): 2 EHT pol+ cells with EHT cell 1 more advanced in its emergence than EHT cell 2; mKate2-podxl2 expressed by EHT cell 1 is enriched at the apical pole of the cell, facing the aortic lumen (as is the case for eGFP-podxl2, see ***Figure 1C***). Top panel, right box (with 2X magnification beneath): 3 EHT pol- cells with mKate2-podxl2 localized relatively homogenously between luminal and abluminal membranes. All EHT pol+ and EHT pol- cells express cytosolic eGFP, as seen in top and bottom panels (2 channels images). Note that it is also the case for the he cell (hemogenic cell) in the aortic floor, in the middle of the image, top panel. Bar = 10$\mu$m.

**Figure 1 – video 1. EHT pol+ cells at early timing.** Z-stack (37 planes) corresponding to the time point t = 0 min whose plane 21 is shown ***Figure 1C***.

**Figure 1 – video 2. EHT pol+ cells at late timing.** Z-stack (37 planes) corresponding to the time point t = 80 min whose planes 16 (for eht cell 1) and 22 (for eht cell 2) are shown ***Figure 1 – figure supplement 1B.***

**Figure 1 – video 3. Emergence of EHT pol- cells.** Time-lapse sequence using a *Tg(Kdrl:Gal4;UAS:RFP;4xNR:eGFP-podxl2)* embryo, obtained with spinning disk confocal microscopy (38 timing points, each acquired with 10 min intervals, starting at 55 hpf) and from which the images of the 2 emerging EHT pol- cells shown ***Figure 1D*** were extracted.

**Figure 2 – figure supplement 1. HE cells are not polarized at 30 hpf.** *Tg(Kdrl:Gal4;UAS:RFP;4xNR:eGFP-podxl2)* 30 hpf embryo imaged using spinning disk confocal microscopy. **Top 2 panels**: green (eGFP-podxl2) and red (soluble RFP, in magenta) channels. White arrows point at 2 individual EHT cells. Note that HE cell 1 protrudes long filopodia, some of which inside the aortic lumen (live imaging shows that they are moving along the blood flow, data not shown). **Bottom 2 panels**: 1.625X magnification of EHT cell 1 and 2 in top panels. Green channel (eGFP-podxl2) only. Green arrows point at very large intracellular vesicles, the largest reaching approximately 30$\mu$m. Asterisks mark the cytoplasm. Bar: 100$\mu$m.

**Figure 2 – figure supplement 2. Evolution of non-polarized HE cells throughout emergence**. *Tg(Kdrl:Gal4;UAS:RFP;4xNR:eGFP-podxl2)* embryo imaged using spinning disk confocal microscopy. Images were obtained from discontinuous time-lapse sequences covering a period of 13 hours (from 35 – to – 48 hpf). Successive phases of the evolution of HE cells are visible, from a non-polarized status (with the accumulation of Podxl2-containing intra-cytosolic vesicles as well as cell division) to post-emergence EHT cells remaining beneath the aortic floor. The top panel is a z-projection of 69 consecutive z-sections interspaced by 0.3$\mu$m; green (eGFP-podxl2) and red (soluble RFP, in magenta) channels are shown; 4 individual HE cells (1 – 4, green arrows)) are marked, with cells 1 and 4 more advanced in the process of emergence. All the other panels are z-sections focused on the aortic floor, allowing visualizing the progression of the EHT, in particular at 35 hpf (t = 0), a timing point at which the separation between the luminal and basal membranes are equally labelled with eGFP-podxl2, attesting for the absence of apicobasal polarity (see cells 1 and 4, see also cell 1 at t = 2.5 hours (hrs) and cells 3' and 3" at t = 5 hrs). Note, at t = 2.5 and 5.0 hrs, the presence of intracytoplasmic eGFP-podxl2 labelled vesicles inside HE cells, suggesting vesicular transport (white arrowheads, to be compared with images ***Figure 2 – figure supplement 1***, at 30 hpf). At 48 hpf (bottom panel), HE cells have emerged and remain, for some of them, in close contact with the aortic floor (after having performed mitosis (notably for HE cell 2 (the division is visualized at 5hrs) and for HE cell 3, each one giving rise to cells 2' - 2" and 3' - 3" respectively, all containing residual eGFP-podxl2 containing membranes as EHT signature). Bar = 80$\mu$m.

**Figure 3 – figure supplement 1. Expression of dt-runx1 triggers the accumulation of myb+ cells in the aortic floor. A**, representative image (Imaris 3D-rendering) of RNA-scope in situ hybridization for the hematopoietic marker myb in 48 - 50hpf *Tg(Kdrl:eGFP*) control embryos (**left**) and 48 - 50hpf *Tg(Kdrl:Gal4;UAS:RFP;4xNR:dt-runx1-eGFP)* mutant embryos (**right**). **Top row**: 3D rendering without segmentation. The aorta is outlined with the white dashed lines. **Center top row**: Magnification of aortic segments (outlined with white boxes on the top rows). Cyan arrows: hemogenic cells (comprising characterized elongated hemogenic cells as well as EHT undergoing cells). Magenta arrows: hematopoietic cells in the sub-aortic space. **Center bottom row**: GFP+ cells segmentation (white cellular outline) and RNAscope signal segmentation (magenta spots). **Bottom row:** Manual cell classification post segmentation. Aortic roof cells (virtually all negative for myb) are displayed in green, hemogenic and EHT cells in cyan and sub-aortic hematopoietic cells in magenta (all expressing myb). Scale bars: 10$\mu$m. **B,** quantitative analysis of the dt-runx1 expressing mutant phenotype. **Left**: endothelial roof cell count (green segmented cells on panel **A**). **Right**: hemogenic/EHT cell count (cyan segmented cells on panel **A**). **C**, myb positive cell count in roof endothelial cells (**left**, corresponding to green cells in panel **A**), hemogenic/EHT cells (**middle**, corresponding to cyan cells in panel **A**) and hematopoietic cells in the sub-aortic region (**right**, corresponding to magenta cells in panel **A**). **D**, number of segmented RNAscope myb dots per segmented roof cells (**left** panel) and hemogenic/EHT cells (**right** panel). For panels **B**, **C** and **D**, statistical comparisons have been performed using two-sided unpaired Wilcoxon tests, all p-values are displayed. Analysis was carried out on n = 4 control embryos and n = 3 mutant embryos, with 2 aortic segments per embryo.

**Figure 3 – figure supplement 2. Phenotypic analysis of dt-runx1 expressing mutants: evidence for apicobasal polarity of hemogenic cells.** 30 - 32 hpf and 48 - 55hpf embryos obtained from *outcrossed Tg(Kdrl:mKate2-podxl2) X Tg(kdrl:Gal4;UAS:RFP;4xNR:dt-runx1-eGFP)* fishes were imaged in the trunk (AGM) region using spinning disk confocal microscopy (single z-planes). **A,** two aortic segments imaged in control sibling embryos (not expressing the dt-runx1 mutant, hence negative for the cleaved eGFP). Typical HE cells are outlined in the white boxes (with the luminal and abluminal membranes clearly separated from each other owing to reduction of their surface area (in comparison to flat aortic cells) and elongation in the antero-posterior axis). Note the absence of enrichment of mKate2-podxl2 in luminal membranes in comparison to basal membranes (magenta arrows). **B,** two aortic segments in dt-runx1 expressing embryos. Images are depicting typical HE cells (in boxes) from hemogenic regions (as in **A**). Note the enrichment of mKate2-podxl2 in apical/luminal membranes (magenta arrows) in comparison to basal membranes (green arrows). Green: cytosolic eGFP released from the cleavage of dt-runx1-eGFP. Note that because of mosaicism, he cell 2 does not express mkate2-podxl2. Scale bars: 10$\mu$m. **C,** evolution of an EHT cell extracted from a 7 hrs time-lapse sequence (0 to 420 min, starting from around 40hpf) and showing significant changes in its morphology throughout emergence. Note the sub-luminal and cytosolic localization of pools of Podxl2 (notably at t = 30 min, magenta arrows) suggesting enhanced trafficking of the protein and relative instability of apical polarity, consistently with apparent fluctuation of the luminal membrane surface contacting the aortic lumen (green arrows, in particular at t = 240 min). At t = 420 min, the cell has emerged and Podxl2 containing membranes remain in close contact with the membrane contacting the aortic floor. Scale bars: 20$\mu$m.

**Figure 3 – figure supplement 3. Phenotypic analysis of dt-runx1 expressing mutants: expansion of the thymus. A**, 3D-rendering of confocal spinning disk sections of the thymus region of 5 dpf *Tg(kdrl:gal4;UAS:RFP)* (left, control) and incrossed *Tg(kdrl:Gal4;UAS:RFP;4xNR:dt-runx1-eGFP)* (right, dt-runx1) larva. Thymic cells have been individually segmented. Bar: 10$\mu$m. **B**, morphometric analysis of the thymus. **Left**: thymic volume (in $\mu m^3$, corresponding to the sum of all thymic cell volumes). Center: cell count. Right: thymic cells volume (in $\mu m^3$). Statistical comparisons have been performed using two-sided unpaired Wilcoxon tests, all p-values are displayed. Analysis was carried out on n = 5 control embryos and n = 5 mutant embryos.

**Figure 3 – video 1. Cellular expansion in the thymus of a dt-runx1 embryo compared to control.** 3D-rendering of confocal spinning disk sections of the thymus region of 5 dpf *Tg(kdrl:gal4;UAS:RFP)* (left, control) and incrossed *Tg(kdrl:Gal4;UAS:RFP;4xNR:dt-runx1-eGFP)* (right, dt-runx1) larva. Thymic cells have been individually segmented. Scale bar: 10$\mu$m.

**Figure 4 – figure supplement 1. Expression levels of Pard3 isoforms in FACS-sorted endothelial cells. A,** experimental strategy to isolate endothelial cells for subsequent gene expression analyses by qRT-PCR. 48hpf *Tg(Kdrl:Gal4;UAS:RFP)* whole embryos and trunk dissected regions (delimited by two magenta arrows) were treated to dissociate cells and populations of interest were sorted by FACS. n = 2 experiments were performed for the whole embryo RFP+ cell population (**left**) and n = 3 experiments were performed for the trunk RFP+ cell population (**right**). Subsequently, RNA was isolated from 20 000 negative cells (RFP-, black) and from 6 500 to 9 142 cells for the RFP expressing population (RFP+, magenta). The relative proportions of RFP+ and RFP- cells in the two conditions are displayed (table). **B**, **C**, qRT-PCR analysis of gene expression levels of the hematopoietic markers myb and runx1 (**B**) and of the 4 Pard3 isoforms (Pard3aa, ab, ba and bb, **C**) in vascular and non-vascular cells from 48 - 50hpf whole embryos and dissected trunks. Graphs show the measured mean fold changes relative to ef1α. Statistical tests (two sided unpaired two samples Wilcoxon test, all p-values are displayed) were only performed to compare expression levels in RFP+ and RFP- cells isolated from trunk regions (three independent experiments).

**Figure 4 – figure supplement 2. mRNA localization of Pard3 isoforms using RNAscope.** **A, B, C,** representative images (Imaris 3D-rendering) of RNAscope *in situ* hybridizations for Pard3aa, Pard3ba and Pard3ab in 48 - 50hpf *Tg(Kdrl:eGFP*) control embryos. Images show the whole pattern of expression of Pard3aa, Pard3ba and Pard3ab mRNAs in the trunk region (z-projections). Anatomical regions are outlined with white dashed lines, and roughly annotated based on location and with the support of transmitted light images (not shown). The aorta is outlined with magenta dashed lines. DLAV: dorsal longitudinal anastomotic vessels, SC: spinal cord, NT: notochord, SAS: sub-aortic space, PCV: posterior cardinal vein, PD: pronephric duct. Scale bars: 10$\mu$m. **D**, representative images (Imaris 3D-rendering) of RNAscope *in situ* hybridizations for Pard3aa, Pard3ba and Pard3ab from trunks narrowed down to aortic and sub-aortic regions of 48 - 50hpf *Tg(Kdrl:eGFP*) embryos. The aorta is outlined with white dashed lines. The RNAscope signals were segmented into spots and classified based on their localization: in aortic endothelial and hemogenic cells (magenta spots) or in extra-aortic tissues (grey spots). Scale bars: 10$\mu$m.

**Figure 4 – figure supplement 3. Expression of Pard3ba is upregulated by dt-Runx1. A,** experimental strategy to isolate endothelial cells from *Tg(Kdrl:Gal4;UAS:RFP)* and *Tg(Kdrl:Gal4;UAS:RFP;4xNR:dt-runx1-eGFP)* embryos for subsequent gene expression analyses by qRT-PCR. 48hpf control (**left**) and dt-runx1 expressing mutant embryos (**right**) were dissected to isolate trunk regions (delimited by two magenta arrows) that were subsequently treated to dissociate cells and populations of interest were sorted by FACS. n = 3 experiments were performed. RNA was isolated from 1 900 to 3 200 cells for the dt-runx1 mutant (green population) and from 6 500 to 9 142 cells for control RFP expressing cells (magenta population). **B**, representative images (Imaris 3D-rendering) of RNAscope *in situ* hybridizations for Pard3ba in 48 - 50hpf *Tg(Kdrl:eGFP*) control embryos (**left**) and dt-runx1 mutant embryos *Tg(Kdrl:Gal4;UAS:RFP;4xNR:dt-runx1-eGFP)* (**right**). The aorta is outlined with the white dashed lines. The RNAscope signals were segmented into spots and classified based on their localization: in aortic endothelial and hemogenic cells (magenta spots) or in extra-aortic tissues (grey spots). Scale bars: 10$\mu$m. **C,** hemogenic/EHT cell count per aorta segment, separated based on their pard3ba expression (with, from left to right, 0 to 4 Par3ba spots). Statistical tests: two sided unpaired two samples Wilcoxon test, all p-values are displayed. Analysis was carried out on n = 5 control embryos and n = 6 mutant embryos, 2 aortic segments per embryo.

**Figure 4 – figure supplement 4. Expression of Pard3aa and Pard3ab are insensitive to dt-Runx1. A, B,** representative images (Imaris 3D-rendering) of RNAscope *in situ* hybridizations for Pard3aa and Pard3ab in trunk regions of 48 - 50hpf *Tg(Kdrl:eGFP*) control embryos and *Tg(Kdrl:Gal4;UAS:RFP;4xNR:dt-runx1-eGFP)* mutant embryos. The aorta is outlined with the white dashed lines. The RNAscope signals were segmented into spots; only the spots localized in the dorsal aorta are represented (magenta spots). Scale bars: 10$\mu$m. **C**, Pard3aa (**left**) and Pard3ab (**right**) spots count per aortic segment. Statistical tests: two sided unpaired two samples Wilcoxon test, all p-values are displayed. Analysis was carried out on n = 3 control embryos and n = 3 mutant embryos.

**Figure 5 – figure supplement 1. Model depicting the evolution of junctional interfaces and of the differential mobility of antero-posterior junctional complexes for EHT pol+ and EHT pol- cell emergence types. Top panel:** EHT pol+ cell whose emergence depends on the constriction of circumferential actomyosin (orange, see Lancino et al. 2018) and the reduction of the membrane interface contacting endothelial neighbors shrinking along the longitudinal axis (horizontal double arrows) and in the 2D plane (endothelial neighbors are not depicted and are embedded in the blue X-Y plane). Presumably, the reduction of membrane interfaces relies on consumption via endocytosis. Green ovals: junctional complexes reinforced at antero-posterior poles. Green line: apical/luminal membrane with typical inward bending. **Bottom panel**: EHT pol- cell whose emergence depends on the dynamics of adhesion pools that move synchronously in 3D (X, Y, Z), both at antero-posterior poles (blue ovals) and at lateral sides of the emerging cell contacting endothelial neighbors (adjoining endothelial cells are not depicted and are embedded in the blue X-Y plane). Presumably, this type of emergence in which endothelial cells crawl over the EHT cell (curved arrows) involves, for the latter, a partial retrograde endocytic recycling of junctional complexes (opposite to the direction of emergence). Note that nuclei are not drawn at scale.

**Figure 5 – figure supplement 2. JAM2a and JAM3b expression and localization in diverse embryonic tissues.** Localization of transiently expressed eGFP-Jam2a (**A-D**) and eGFP-Jam3b (**E-G**) in 52 hpf embryos, using spinning disk confocal microscopy. Plasmid constructs were expressed under the control of the heat shock *Hsp70* promoter. For both constructs, expression was induced approximately 6 hrs before imaging, by 1hour balneation in 39°C embryo medium. All images are maximum z-projections. **A, E,** ependymal cells. White and red arrows point at reinforcement of eGFP-Jam2a and eGFP-Jam3b at apical intercellular junctions and at basolateral membranes, respectively. **B,** pronephric tubule cells. White arrows point at the reinforcement of the eGFP-Jam2a signal at apical sides of membranes of polarized cells constituting pronephric tubules. The red arrow points at baso-lateral localization of eGFP-Jam2a. **C, F,** skin epithelial cells. White arrows point at the localization of eGFP-Jam2a and eGFP-Jam3b at lateral junctional interfaces between two neighboring cells. Red arrows point at tri-cellular junctions at which the density of eGFP-Jam2a and eGFP-Jam3b is reinforced. **D,** skin epithelial cells. White arrows point at membrane protrusions. **G,** Striated muscle cells. Red arrows point at T-tubules (invagination of the sarcolemmal membranes); white arrows point at the plasma membranes of myofibrils. Bars: 20$\mu$m.

**Figure 6 – figure supplement 1. Examples of junctional contacts targeted by FRAP in the aortic landscape.** After performing z-stack acquisitions in trunk regions followed by 2D-deployment of aortic segments (**A-C**), 4 different *Tg(kdrl:eGFP-Jam3b; kdrl:nls-mKate2)* 48-55 hpf embryos were illuminated for FRAP. Panels (**A)** and (**B)** are the entire segments from which cropped images of EHT pol+ and EHT pol- cells (black asterisks) and surroundings were extracted to build the panels A and B of ***Figure 6***. Panels (**C**) and (**D**) are from 2 other embryos. The 2D-cartographies allow visualizing the junctions that were selected for FRAP, with the black arrows pointing at junctional interfaces between EHT and endothelial cells (all are in the area of tri-junctions) and the green arrows pointing at bi-junctions (Bj) or tri-junctions between endothelial cells (green asterisks). Bar = 20$\mu$m.

**Figure 7 – figure supplement 1. Searching for PDZ-domain containing RhoGEFs potentially involved in the EHT. A,** cartoons representing the domains composing the 9 PDZ domain-containing RhoGEFs that were investigated in this study (all these RhoGEFs are encoded by different genes in the zebrafish; protein respective length is not drawn at scale). **B,** results obtained from 3 independent qRT-PCR experiments and performed on material extracted from trunk regions of 35 and 48 hpf embryos obtained from incrossed *Tg(kdrl:Gal4;UAS:RFP;4xNR:dt-runx1-eGFP)* adult fishes. Statistical tests: two sided unpaired two samples Wilcoxon test, with p-values of significant differences. GEF: guanine nucleotide-exchange factor; PDZ: postsynaptic density protein of 95 kDa, Discs large and Zona occludens-1; PH: Plekstrin homology; RGS: regulator of G-protein signaling; DH: Dbl (diffuse B-cell lymphoma) homology; RBD: Ras-binding domain; DEP: Dishevelled Egl-10 and

Plekstrin; InsPx4-PTPase: PtdIns(1,3,4,5)P4 phosphatidylinositol phosphatase; TIAM: T-cell-lymphoma invasion and metastasis; LARG: leukemia-associated Rho guanine-nucleotide exchange factor; PRex: PtdIns(3,4,5)P3-dependant Rac exchanger-1 and 2.

**Figure 7 – figure supplement 2.** Whole mount in situ hybridizations (WISH) performed on 30-32 hpf and 48-50 hpf embryos, with probes specific for all 9 PDZ-domain containing RhoGEFs that were investigated in this study. Note that all RhoGEFs are detected in the dorsal aorta of the trunk region (white arrowheads).

**Figure 7 – figure supplement 3. A N-terminal fragment of ArhGEF11/PDZ-RhoGEF localizes at junctional membranes with enrichment at antero-posterior sites of EHT cells. A,** domains of ArhGEF11, including its actin binding site (L/IIxxFE) and the position of exon 38 encoded peptide. The bottom cartoon represents a protein deleted from its DH-PH domains and C-terminus that are replaced by eGFP (PDZ-PRD-RGS-eGFP) to follow the localization of the truncated fusion protein in expressing cells. **B**, localization of the ArhGEF11 PDZ-PRD-RGS-eGFP fusion protein after transient expression in the aorta (expression was obtained after injection of the plasmid, at the one cell stage, in the *Tg(Kdrl:Gal4;UAS:RFP)* fish line). Confocal images (z-projection) extracted from a time-lapse sequence (timing points t = 0 and t = 77 min) and showing either the eGFP channel only (left), or the merge between the green and the red channels (RFP labels the cytoplasm). Note the localization of PDZ-PRD-RGS-eGFP at membrane interfaces at early time point of the emergence and at the rim of an emerging cell (white arrowheads, second cell from the right) and its enrichment at antero-posterior sites of an EHT pol+ cell proceeding throughout emergence (right cell, at t = 0 and t = 77 min and bottom cartoons, green arrows point at localization/concentration of PDZ-PRD-RGS-eGFP ).

**Figure 7 – figure supplement 4. MO and CRISPR approaches to investigate the function of ArhGEF11/PDZ-RhoGEF in the EHT. A**, splicing MO interfering with integration of exon 38. **Top panel**: cartoon representing exons/introns (not drawn at scale) composing the 3-prime region of the gene encoding for ArhGEF11, with the position of the splicing MO. **Middle panel**: agarose gel showing the 2 alternative mRNAs encoding for ArhGEF11 in control animals (left track, control) and after injection of the MO at the one cell stage (right track, +MO at 2 and 5ng). **Bottom panel**: ArhGEF11 DNA sequences obtained after RT-PCR, cloning and sequencing for 1 control and 2 +MO clones (+MO1, +MO3). **B**, morphologies of control and +MO injected 48 hpf embryos. Note the absence of malformations. **C**, **top panel**: ArhGEF11 wild type and CRISPR-mediated 7bp deletion in the 3-prime region of exon 38 leading to a frame-shift in the ORF and a downstream premature stop codon; **middle panel**: CRISPR del/C-ter nucleotide/aa sequence and wild type nucleotide/aa sequence covering the extreme C-ter of the full-length protein; **bottom panel**: CLUSTAL 2.1 multiple sequence alignment of mouse and zebrafish C-termini highlighting the sequences of spliced variants of potentially similar activity in the regulation of ArhGEF11 activity on RhoA. Accession numbers: NP_001003912.1 and NP_001027010.1 for the mouse and Danio rerio sequence, respectively. **D,** morphologies of control and homozygous ArhGEF11$^{CRISPR\text{-}Cterdel+/+}$ 48 hpf embryos (*(Kdrl:eGFP-Jam3b; kdrl:*ArhGEF11$^{CRISPR\text{-}Cterdel+/+}$) embryos). Note the edema in the cardiac region (35 hpf embryos also exhibited retardation in blood circulation).

**Figure 7 – figure supplement 5. Supplementary data on the ArhGEF11/PDZ-RhoGEF exon 38 splicing morpholino phenotype.** *Tg(kdrl:eGFP-Jam2a; kdrl:nls-mKate2)* control (**A, C, E**) or MO-injected (**B, D, F**) embryos were imaged using spinning disk confocal microscopy (48-55 hpf time-window). Aorta segments (330 $\mu$m each) were imaged in the trunk region (AGM). The 2D-cartographies with delimited cellular contours in panels **A** and **B** are presented ***Figure 7A*** with the corresponding un-modified 2D-cartographies shown on top for unmasking of contours. Panels **A** and **C** illustrate 2 different segments from the same embryo. Panels **B,**

**D, F** illustrate segments from 3 different embryos. (a-f) Maximum z-projections of merged nls-mkate2 and eGFP-Jam2a signals for control (a, c, e) and for ArhGEF11 exon 38 splicing morpholino (b, d, f) conditions. For panels (a) and (b), maximum z-projections of the eGFP-Jam2a signal only are also shown. (a'-f') Single z-plane images of merged nls-mkate2 and eGFP-Jam2a signals for control (a', c', e') and for ArhGEF11 exon 38 splicing morpholino (b', d', f') conditions. For **e'**, the image is a composition of 2 different z-planes from the same field (the boundaries are marked with white ticks). In right margins, magenta and green arrowheads designate the aortic floor and roof, respectively. (a''-f'') 2D-cartographies (bottom, with delineated cellular contours) obtained from eGFP-Jam2a signals for control (a'', c'', e'') and for the ArhGEF11 exon 38 splicing morpholino (b'', d'', f'') conditions, respectively. Cell contours are delineated either in blue (endothelial cells), yellow (hemogenic cells), red (morphologically characterized EHT cells, red arrows), and small cells delineated by cyan boxes (morphologically uncharacterized EHT cells and putative post-mitotic cells remaining as pairs). Cellular contours have been semi-automatically segmented along the cellular interfaces labelled with eGFP-Jam3b (see ***Methods***). White and black arrows designate hemogenic cells with their nucleus visible on the z-section. Analyses were performed on 2 x control non-injected embryos and 3 x embryos injected at the one-cell stage with the ArhGEF11 exon 38 splicing MO. Bars = 20$\mu$m.

**Figure 7 – figure supplement 6. Supplementary data on the ArhGEF11/PDZ-RhoGEF CRISPR C-ter deletion phenotype.** *(Kdrl:eGFP-Jam3b; kdrl:*ArhGEF11$^{\text{CRISPR-Cterdel+/+}}$) homozygous ArhGEF11 C-ter deletion mutants (CRISPR ArhGEF11: panels **B, D, F**) and control siblings (Control: panels **A, C, E**) were imaged using spinning disk confocal microscopy (48-55hpf time-window). Note that the genetic background of the CRISPR fish line is (Kdrl:GAL4; UAS:RFP) thus allowing the red cytosolic staining of aortic and hemogenic cells. Aorta segments (330 $\mu$m each) were imaged in the trunk region. The 2D-cartographies with delimited cellular contours in panels **A** and **B** are presented ***Figure 7B*** with the corresponding un-modified 2D-cartographies shown on top for unmasking of contours. Panels **A, C** and **D, F** illustrate 2 different segments from the same embryo. (a-f) Maximum z-projections of merged nls-mkate2 and eGFP-Jam3b signals for control (a, c, e) and for the CRISPR mutant (b, d, f) conditions. For panels (a) and (b), maximum z-projections of the eGFP-Jam2a signal only are also shown. (a'-f') Single z-plane images of merged nls-mkate2 and eGFP-Jam3b signals for control (a', c', e') and for ArhGEF11 exon 38 splicing morpholino (b', d', f') conditions. For b' and d', the image is a composition of 2 different z-planes from the same field (the boundaries are marked with white ticks). In right margins, magenta and green arrowheads designate the aortic floor and roof, respectively. (a''-f'') 2D-cartographies (bottom, with delineated cellular contours) obtained from eGFP-Jam3b signals for control (a'', c'', e'') and for the CRIPR mutant (b'', d'', f'') conditions, respectively.

Cell contours are delineated either in blue (endothelial cells), yellow (hemogenic cells), red (morphologically characterized EHT cells, red arrows), and small cells delineated by cyan boxes (morphologically uncharacterized EHT cells). Cellular contours have been semi-automatically segmented along the cellular interfaces labelled with eGFP-Jam3b (see ***Methods***). White and black arrows designate hemogenic cells with their nucleus visible on the z-section. Analyses were performed on 2 x wild-type siblings for controls and 2 x mutant embryos whose mutation was confirmed by DNA sequencing. Bars = 20$\mu$m.

**Figure 7 — Figure supplement 7. MO interference with ArhGEF11 exon38 splicing leads to the accumulation of CD41 positive cells in the aortic floor and negatively impacts on the generation of hematopoietic precursors. A, B,** representative images of AGM region of double transgenic *Tg(CD41:eGFP; Kdrl:nls-mKate2)* embryos, injected with control (**A**) or ArhGEF11 exon38 splicing morpholino (**B**) at 48 - 55hpf. The two top panels show maximum z-projection of aortic segments (top panels with the CD41:eGFP signal in green and the Kdrl:nls-mKate2 signal in magenta, bottom panel only with the CD41:eGFP signal in white).

The three bottom panels show single z-planes extracted from the above z-projections. Automatically segmented GFP positives cells are outlined in white. Magenta arrows point at double positives (GFPlow/kdrl+ cells), corresponding to hemogenic/EHT cells. Blue arrows point at single positive (GFPlow/kdrl- cells) HSPC precursors. Green arrows point at circulating thrombocytes that were automatically eliminated from the segmentation. The asterisk labels a non-moving thrombocyte (GFPhigh/kdrl- cells). The aortic floor is outlined by the white dashed line. Scale bars: 10$\mu$m. **C**, GFP and mKate2 fluorescence intensity was automatically extracted for each segmented cell and normalized to background. Vertical and horizontal dashed lines on the plot and on the zoomed plot represent the threshold selected for the classification of mKate2+ versus mKate2- (1.05, horizontal line) and eGFPhigh versus eGFPlow (1.4, vertical line). **D**, hemogenic/EHT cell count. **E**, HSPC precursors cell count. For **D** and **E**: Statistical tests: two sided unpaired two samples Wilcoxon test, all p-values are displayed. Analysis was carried out on n = 8 control and n = 6 mutant embryos.

#### Supplementary files

**Supplementary file 1**: Materials and Methods - Supplementary Table

**Figure 1 – video 1. EHT pol+ cells at early timing.** Z-stack (37 planes) corresponding to the time point t = 0 min whose plane 21 is shown ***Figure 1C***.

**Figure 1 – video 2. EHT pol+ cells at late timing.** Z-stack (37 planes) corresponding to the time point t = 80 min whose planes 16 (for eht cell 1) and 22 (for eht cell 2) are shown ***Figure 1 – figure supplement 1B.***

**Figure 1 – video 3. Emergence of EHT pol- cells.** Time-lapse sequence using a *Tg(Kdrl:Gal4;UAS:RFP;4xNR:eGFP-Podxl2)* embryo, obtained with spinning disk confocal microscopy (38 timing points, each acquired with 10 min intervals, starting at 55 hpf) and from which the images of the 2 emerging EHT pol- cells shown Figure 1D were extracted.

**Figure 3 – video 1. Cellular expansion in the thymus of a dt-runx1 embryo compared to control.** 3D-rendering of confocal spinning disk sections of the thymus region of 5 dpf *Tg(kdrl:gal4;UAS:RFP)* (left, control) and incrossed *Tg(kdrl:Gal4;UAS:RFP;4xNR:dt-runx1-eGFP)* (right, dt-runx1) larva. Thymic cells have been individually segmented. Scale bar: 10μm.

| Reagent type (species) or resource | Designation | Source or reference | Identifiers | Additional information |
|---|---|---|---|---|
| recombinant DNA reagent | Oligo for Cloning: 4XNR_CD8-eGFP_fw | This paper | N/A | CCTCGAAGACGCGTGGATCCACCGGTCGCCACCATGGCCTTACCAGTGACCGCCTTGCTCCTGCCGCTGGCCTTGCTGCTCCACGCCGCCAGGCCGAGCCAGTTCCGGGTGTCGGTGAGCAAGGGCGAGGAGCTGTTCACC |
| recombinant DNA reagent | Oligo for Cloning: delPodxl2_eGFP_rev | This paper | N/A | CTGTTTCAGTCAGATATTCGGTCCCGCCTCTAGATCCGGTGGATCCCGGGCCCGCGGTACCGTCGACTGCAGAATTCGAAGCTTGAGCTCGAGATCTGAGTCCGGACTTGTACAGCTCGTCCATGCCGAGAG |
| recombinant DNA reagent | Oligo for Cloning: delPodxl2_fw | This paper | N/A | GCGGGACCGAATATCTGACTGAAACAGAGCTTCATGACTCACAGGAGGCCGAACAGG |
| recombinant DNA reagent | Oligo for Cloning: Podxl2_pG1-4XNR_rev | This paper | N/A | CGATATCCTCGAGGGTACCGAATTCTTAAAGGTGAGTGTCTTCCTCTTGATTG |
| recombinant DNA reagent | Oligo for Cloning: pG1-flk1_CD8_mKate2_fw | This paper | N/A | GACGTTATTTTAACAGACAAGGGCGGTCGCCACCATGGCCTTACCAGTGACCGCCTTGCTCCTGCCGCTGGCCTTGCTGCTCCACGCCGCCAGGCCGAGCCAGTTCCGGGTGTCGGTGAGCGAGCTGATTAAGGAGAACATG |
| recombinant DNA reagent | Oligo for Cloning: delPodxL2-mKate2_rev | This paper | N/A | TGTTTCAGTCAGATATTCGGTCCCGCCTCTGTGCCCCAGTTTGCTAGGGAGGTC |
| recombinant DNA reagent | Oligo for Cloning: mKate2-delPodxL2_fw | This paper | N/A | GACCTCCCTAGCAAACTGGGGCACAGAGGCGGGACCGAATATCTGACTGAAACA |
| recombinant DNA reagent | Oligo for Cloning: Podxl2_PG1-flk1_rev | This paper | N/A | GCCGCGGATCCGATATCCTCGAGGGTACCGAATTCTTAAAGGTGAGTGTCTTCCTCTTGATTGTCTTCCTCGTC |
| recombinant DNA reagent | Oligo for Cloning: GB-4xnr-nlsRunx1-fw | This paper | N/A | TCGAAGACGCGTGGATCCACCGGTCGCCACCATGGTTCCAAAGAAGAAG |
| recombinant DNA reagent | Oligo for Cloning: GB-eGFP-T2A-2xHA-delRunx1rev | This paper | N/A | CCCGGTGAACAGCTCCTCGCCCTTGCTCACCATAGGTCCAGGATTCTCCTCCACGTCTCCGCATGTCAGCAGACTTCCTCTTCCCTCTGCGTAATCAGGCACATCATAAGGATAAGCGTAGTCTGGGACGTCGTATGGGTAGGGAATCTGGCTGTGGGCGGGGCTGCCGAAGGG |
| recombinant DNA reagent | Oligo: qPCR pard3ab forward | This paper | N/A | ATTGGAGCGTGGTGGAAAGG |
| recombinant DNA reagent | Oligo: qPCR pard3ab reverse | This paper | N/A | TGAGCTTGCTCAAAGCGAAC |
| recombinant DNA reagent | Oligo: qPCR pard3b forward | This paper | N/A | GAGAATGACTGCATCGTCCG |
| recombinant DNA reagent | Oligo: qPCR pard3b reverse | This paper | N/A | ACATGAAACAGGATTCGCGG |
| recombinant DNA reagent | Oligo: qPCR pard3bb forward | This paper | N/A | GAAGGCCTCTGGGTCTTCAC |
| recombinant DNA reagent | Oligo: qPCR pard3bb reverse | This paper | N/A | AGATGCCTTCCCGTTTGGAG |
| recombinant DNA reagent | Oligo: qPCR pard3ba forward | This paper | N/A | GGCCTCTGGGAATCAAAGTC |
| recombinant DNA reagent | Oligo: qPCR pard3ba reverse | This paper | N/A | TCTCTGAGCGGCTGTTCTTC |
| recombinant DNA reagent | Oligo: qPCR cmyb reverse | N/A | N/A | TCCTGGGGCTCTTGTTTAATCGT |
| recombinant DNA reagent | Oligo: qPCR cmyb forward | N/A | N/A | GCACACCCACTCCATTCAAATCC |
| recombinant DNA reagent | Oligo: qPCR runx1 reverse | N/A | N/A | TGAGCAGTCGTACCCATATCTG |
| recombinant DNA reagent | Oligo: qPCR runx1 forward | N/A | N/A | AGATGGGAGTGATGGGAAGG |
| recombinant DNA reagent | Oligo: qPCR ef1a forward | NA | N/A | CAGCATTATCCAGTCCTTAAGTAGAGTGC |
| recombinant DNA reagent | Oligo: qPCR ef1a reverse | NA | N/A | GCGTCATCAAGAGCGTTGAGAAG |
| recombinant DNA reagent | Oligo for Cloning: PG1-HSP70-tol2-jam2a-eGFP_c1_f | This paper | N/A | CCTGGAATTCGGTACCCTCGAGGATGCCACCATGCTGGTGTGCGTGTCTCTGCTG |
| recombinant DNA reagent | Oligo for Cloning: PG1-HSP70-tol2-jam2a-eGFP_c1_r | This paper | N/A | TTGCTCACCATGGCGCGCCTATCGATCTGCATGTGTGTGTGCTG |
| recombinant DNA reagent | Oligo for Cloning: PG1-HSP70-tol2-jam2a-eGFP_c2_f | This paper | N/A | ACATGCAGATCGATAGGCGCGCCATGGTGAGCAAGGGCGAGGAGC |
| recombinant DNA reagent | Oligo for Cloning: PG1-HSP70-tol2-jam2a-eGFP_c2_r | This paper | N/A | ACGTTCAGGTCGGCGCGCCTCTTGTACAGCTCGTCCATGCCG |
| recombinant DNA reagent | Oligo for Cloning: PG1-HSP70-tol2-jam2a-eGFP_c3_f | This paper | N/A | GCTGTACAAGAGGCGCGCCGACCTGAACGTGGCGGCGGTGG |
| recombinant DNA reagent | Oligo for Cloning: PG1-HSP70-tol2-jam2a-eGFP_c3_r | This paper | N/A | TTATGATCGCGGCCGCGGATCCGATCTACAGCATGAAGGACTGCGTGTGC |
| recombinant DNA reagent | Oligo for Cloning: PG1-HSP70-tol2-jam3b-eGFP_c1_r | This paper | N/A | TTGCTCACCATGGCGCGCCTATAAACTTCCATCATCTGCGGTCC |
| recombinant DNA reagent | Oligo for Cloning: PG1-HSP70-tol2-jam3b-eGFP_c2_f | This paper | N/A | TGATGGAAGTTTATAGGCGCGCCATGGTGAGCAAGGGCGAGGAGC |
| recombinant DNA reagent | Oligo for Cloning: PG1-HSP70-tol2-jam3b-eGFP_c3_r | This paper | N/A | ATTATGATCGCGGCCGCGGATCCGATTCAGATGACAAATGAGGATTTGTGC |
| recombinant DNA reagent | Oligo for Cloning: PG1-HSP70-tol2-jam3b-eGFP_c1_f | This paper | N/A | TTTCCTGGAATTCGGTACCCTCGAGGATGCCACCATGCTGGTGTGCGTGTCTCTGCTGATATTAATTCACAGCGTCCCTGTGTCTCCCGTCACCGTCAGCAGTCGCAATTCTAAACCGTGGGTCAACGAG |
| recombinant DNA reagent | Oligo for Cloning: PG1-HSP70-tol2-jam3b-eGFP_c3_f | This paper | N/A | AGCTGTACAAGAGGCGCGCCGACATCAACATTGCCGGCATCATC |

| Reagent type (species) or resource | Designation | Source or reference | Identifiers | Additional information |
|---|---|---|---|---|
| recombinant DNA reagent | Oligo for Cloning: PG1-HSP70-tol2-jam3b-eGFP_c2_r | This paper | N/A | CAATGTTGATGTCGGCGCGCCTCTTGTACAGCTCGTCCATGCCG |
| recombinant DNA reagent | Oligo: qPCR ArhGEF11_forward | This paper | N/A | ATCACAGAGCAGCGTCCAG |
| recombinant DNA reagent | Oligo: qPCR ArhGEF11_reverse | This paper | N/A | AGTTTGTGGTGATCCGCTTC |
| recombinant DNA reagent | Oligo: qPCR ArhGEF12a_forward | This paper | N/A | AGGATTACGCAGCGTACCTG |
| recombinant DNA reagent | Oligo: qPCR ArhGEF12a_reverse | This paper | N/A | TCAGGCTGATCCCAGAGTCA |
| recombinant DNA reagent | Oligo: qPCR ArhGEF12b_forward | This paper | N/A | TTCTCAAAGCACTGGAGGACC |
| recombinant DNA reagent | Oligo: qPCR ArhGEF12b_reverse | This paper | N/A | CTTGATTTCGGTCTCCTTCAGC |
| recombinant DNA reagent | Oligo: qPCR Prex1_forward | This paper | N/A | TGTGTCGTTGGAGCAAGCTA |
| recombinant DNA reagent | Oligo: qPCR Prex1_reverse | This paper | N/A | GCGGCATCTGGTCCATTACT |
| recombinant DNA reagent | Oligo: qPCR Prex2_forward | This paper | N/A | CGCTAAGTGTCACGTTGGAAC |
| recombinant DNA reagent | Oligo: qPCR Prex2_reverse | This paper | N/A | TGCGATCTCTCACTCCCAGA |
| recombinant DNA reagent | Oligo: qPCR Tiam1a_forward | This paper | N/A | CACGACCCACCATGAACAGA |
| recombinant DNA reagent | Oligo: qPCR Tiam1a_reverse | This paper | N/A | TTCTGTGGGGGTGATTGGTG |
| recombinant DNA reagent | Oligo: qPCR Tiam1b_forward | This paper | N/A | AAGGACTGCAAATCCCAGCA |
| recombinant DNA reagent | Oligo: qPCR Tiam1b_reverse | This paper | N/A | TGTGTCTCGTTCTCTGCACC |
| recombinant DNA reagent | Oligo: qPCR Tiam2a_forward | This paper | N/A | CGTCGAAACATGCAAAGGGT |
| recombinant DNA reagent | Oligo: qPCR Tiam2a_reverse | This paper | N/A | CGGGACTGTCGTTTGAAGGT |
| recombinant DNA reagent | Oligo: qPCR Tiam2b_forward | This paper | N/A | TGTGTCGTTGGAGCAAGCTA |
| recombinant DNA reagent | Oligo: qPCR Tiam2b_reverse | This paper | N/A | GCGGCATCTGGTCCATTACT |
| recombinant DNA reagent | Oligo for WISH probe: ArhGEF11 fw | This paper | N/A | TGAAGAGTGTCCGGATGAAGATCCAGACGC |
| recombinant DNA reagent | Oligo for WISH probe: ArhGEF11 T7rev | This paper | N/A | GAAATTAATACGACTCACTATAGGGTCAGAAACCAGTGGACTGGATTTGAGGTTG |
| recombinant DNA reagent | Oligo for WISH probe: ArhGEF12a fw | This paper | N/A | CGACAGAGACGGTGTGGAGTTAAGCAAGTTC |
| recombinant DNA reagent | Oligo for WISH probe: ArhGEF12a T7rev | This paper | N/A | GAAATTAATACGACTCACTATAGGGTGCTGATCCGACAGGCCTTTGGCGAAG |
| recombinant DNA reagent | Oligo for WISH probe: ArhGEF12b fw | This paper | N/A | ACACTGAACCAATTACTTTAGCGAGCATCC |
| recombinant DNA reagent | Oligo for WISH probe: ArhGEF12b T7rev | This paper | N/A | GAAATTAATACGACTCACTATAGGGATAGCCTTTGGCGTAGCTGGTTATACTTG |
| recombinant DNA reagent | Oligo for WISH probe: PREX1 fw | This paper | N/A | TCGATCAAATCTCTAATTAACAGCCTTCACC |
| recombinant DNA reagent | Oligo for WISH probe: PREX1 T7rev | This paper | N/A | GAAATTAATACGACTCACTATAGGGATTGCAGAGCTCAGAGGACACCAGCAG |
| recombinant DNA reagent | Oligo for WISH probe: PREX2 fw | This paper | N/A | GTGTTAAATCAAGCTCTGAGCACTGAGAG |
| recombinant DNA reagent | Oligo for WISH probe: PREX2 T7rev | This paper | N/A | GAAATTAATACGACTCACTATAGGGACTGCACAGCTCTGAAGCCACGGTGAG |
| recombinant DNA reagent | Oligo for WISH probe: Tiam1a fw | This paper | N/A | CAAGTTCGCAATCTCCCAAATTCCGAGG |
| recombinant DNA reagent | Oligo for WISH probe: Tiam1a T7rev | This paper | N/A | GAAATTAATACGACTCACTATAGGGCAGAAGATGTCACTGTTGCAGCTGT |
| recombinant DNA reagent | Oligo for WISH probe: Tiam1b fw | This paper | N/A | CACGCTGTCCAACACAGATGGAGAGAG |
| recombinant DNA reagent | Oligo for WISH probe: Tiam1b T7rev | This paper | N/A | GAAATTAATACGACTCACTATAGGGCTCTGGCTTTCGTCCTGTGTCTCGTTCTC |
| recombinant DNA reagent | Oligo for WISH probe: Tiam2a fw | This paper | N/A | TCACAAAAGGAGGGCAGGCCTGAGACG |
| recombinant DNA reagent | Oligo for WISH probe: Tiam2a T7rev | This paper | N/A | GAAATTAATACGACTCACTATAGGGTGAAGTTGTCACATTCTTTGGCTCCATGG |
| recombinant DNA reagent | Oligo for WISH probe: Tiam2b fw | This paper | N/A | CTCTTTGGGCCGCATGAGGAAAGACC |
| recombinant DNA reagent | Oligo for WISH probe: Tiam2b T7rev | This paper | N/A | GAAATTAATACGACTCACTATAGGGTGGATGCCCATGAGCGTATCGTAAAAGC |
| recombinant DNA reagent | Oligo for ArhGEF11 exon 38 alternative splicing screen: SM33 forward | This paper | N/A | CTGTCGTCACACCGGCTGATGCAG |
| recombinant DNA reagent | Oligo for ArhGEF11 exon 38 alternative splicing screen: SM30 reverse | This paper | N/A | AGTTTGTGGTGATCCGCTTC |
| recombinant DNA reagent | Oligo for CRISPR PCR screen: SM31 | This paper | N/A | TTTCACTTTCTCTCTGCCTCTTACA |
| recombinant DNA reagent | Oligo for CRISPR PCR screen: SM32bis | This paper | N/A | AACTCTGTCCAGATGATTGAGGAGC |
| recombinant DNA reagent | Oligo for CRISPR PCR screen: SM34 | This paper | N/A | ATAAATGAAGCCCCACCTCCGTCC |
| recombinant DNA reagent | Oligo for CRISPR PCR screen: SM40 | This paper | N/A | ATGAAGCCCCACCTCAGACGATTGGC |

### FIGURE 1 - figure supplement 1

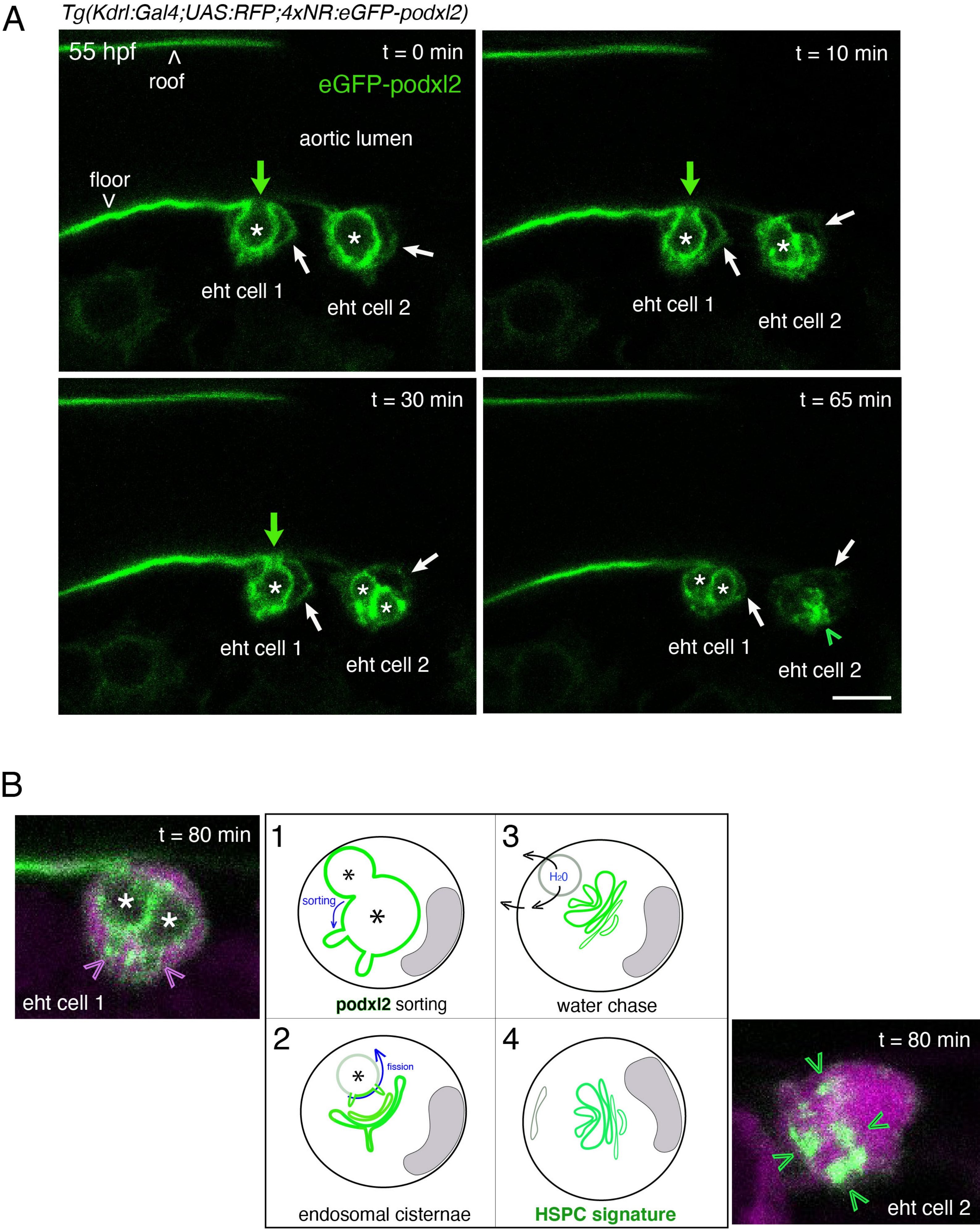

### FIGURE 1 - figure supplement 2

A

*Tg (Kdrl:Jam3b-eGFP; Kdrl:nls-mKate2)*

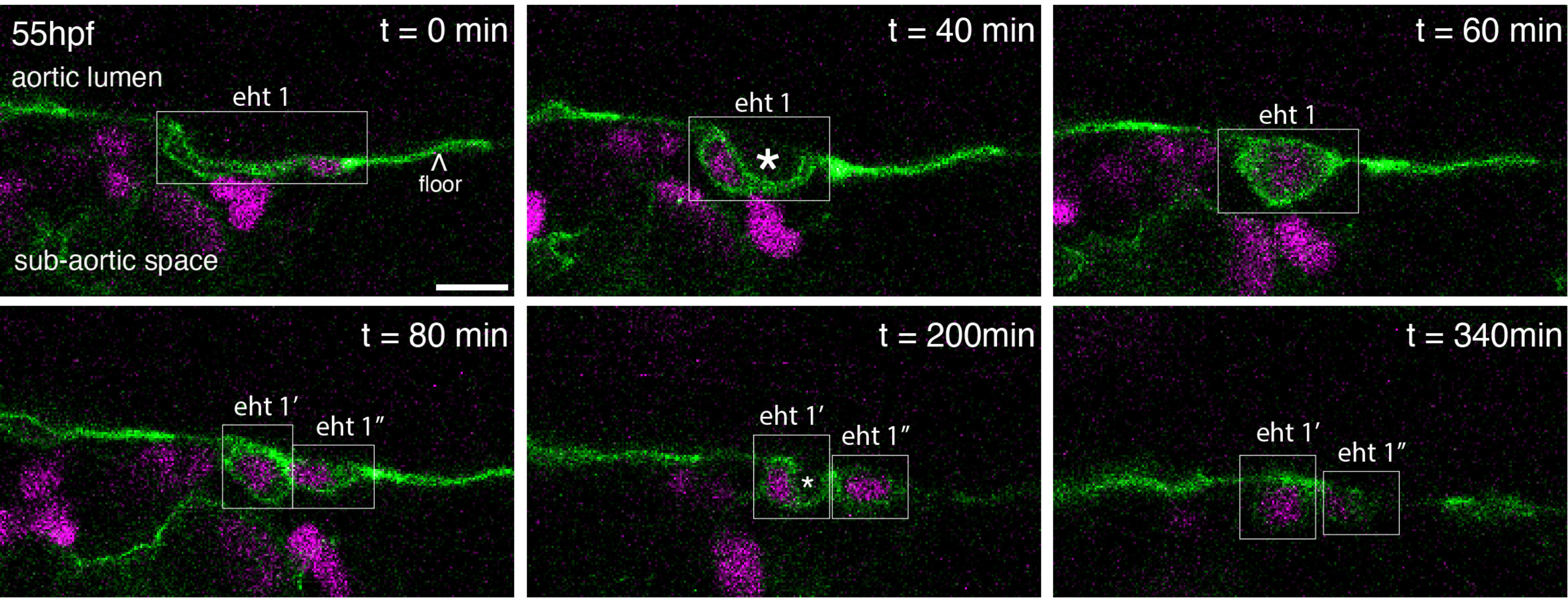


B

*Tg (Kdrl:Gal4;UAS:RFP;4XNR:eGFP-podxl2)*

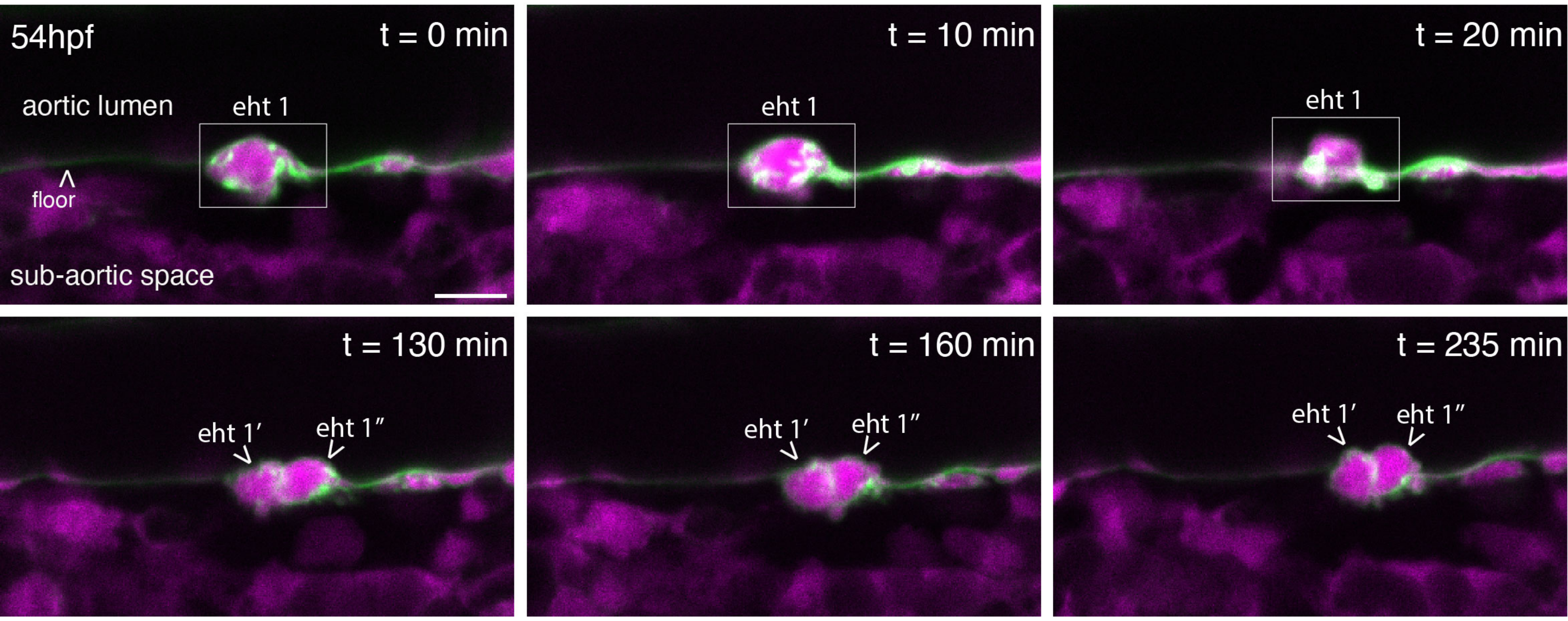

Figure 1 - figure supplement 3

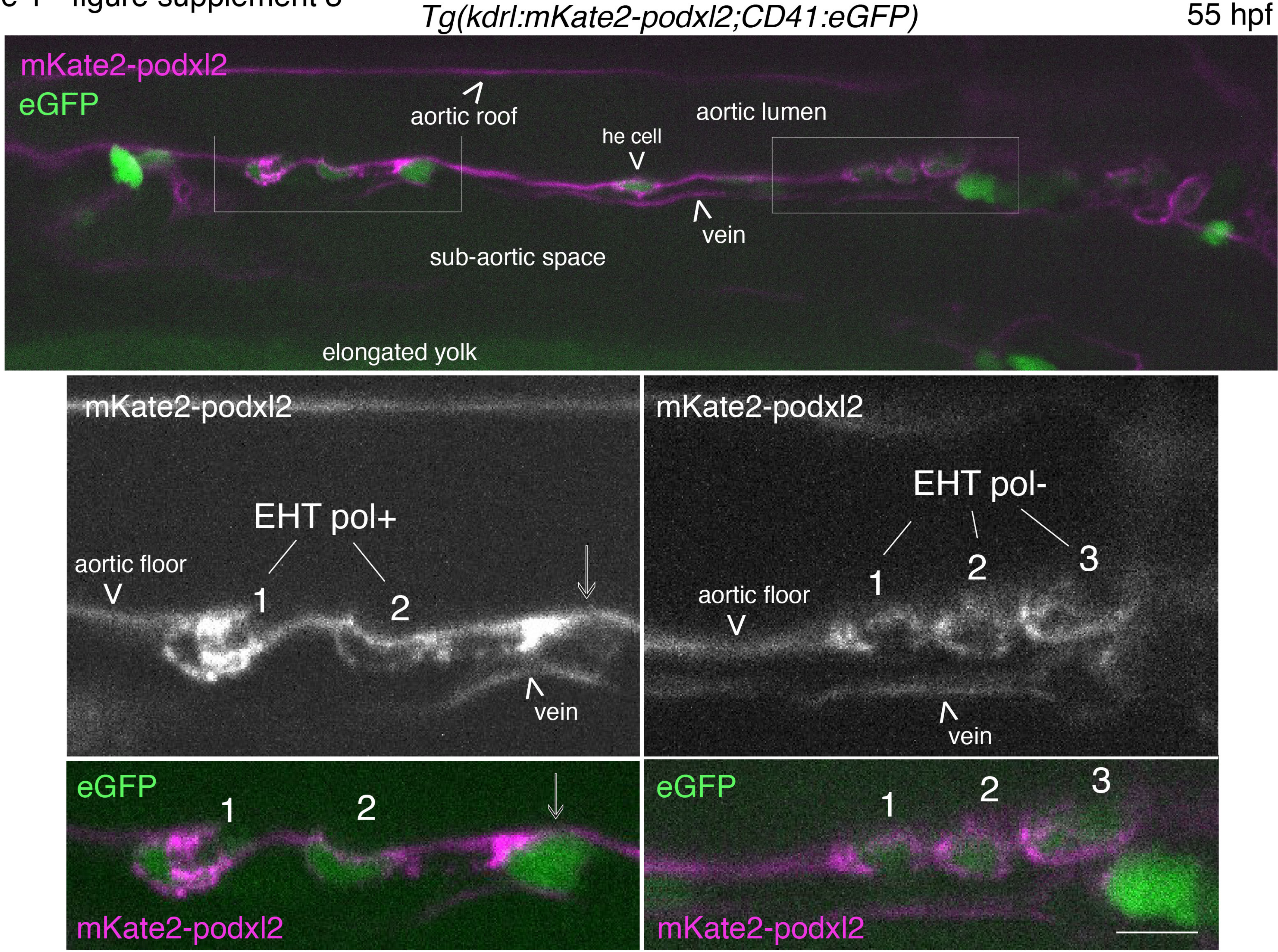

### Figure 2 - figure supplement 1

*Tg(Kdrl:Gal4;UAS:RFP;4xNR:eGFP-podxl2) 30hpf*

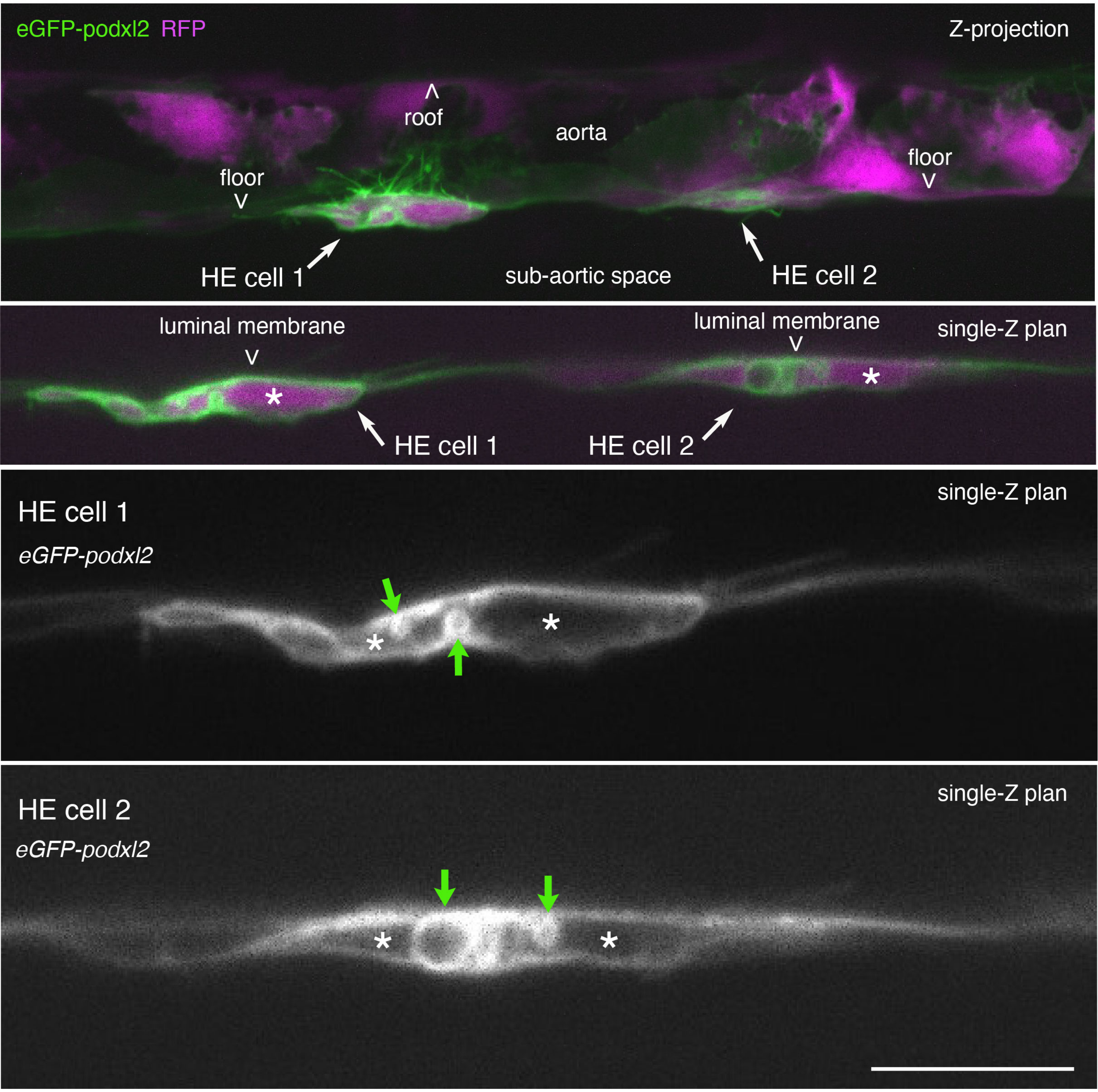

### Figure 2 - figure supplement 2

*Tg(Kdrl:Gal4;UAS:RFP;4xNR:eGFP-podxl2)*

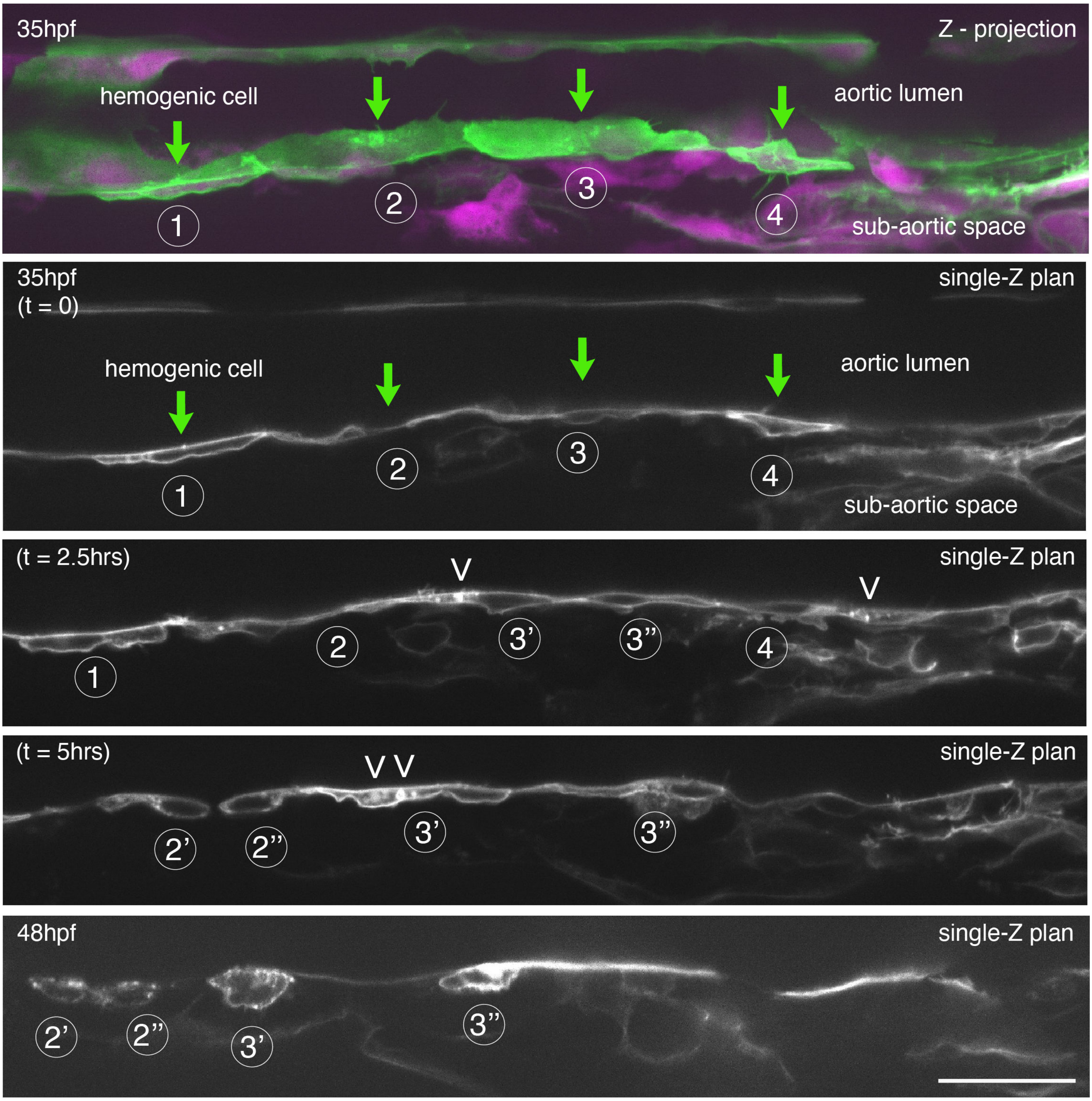

### Figure 3 - figure supplement 1

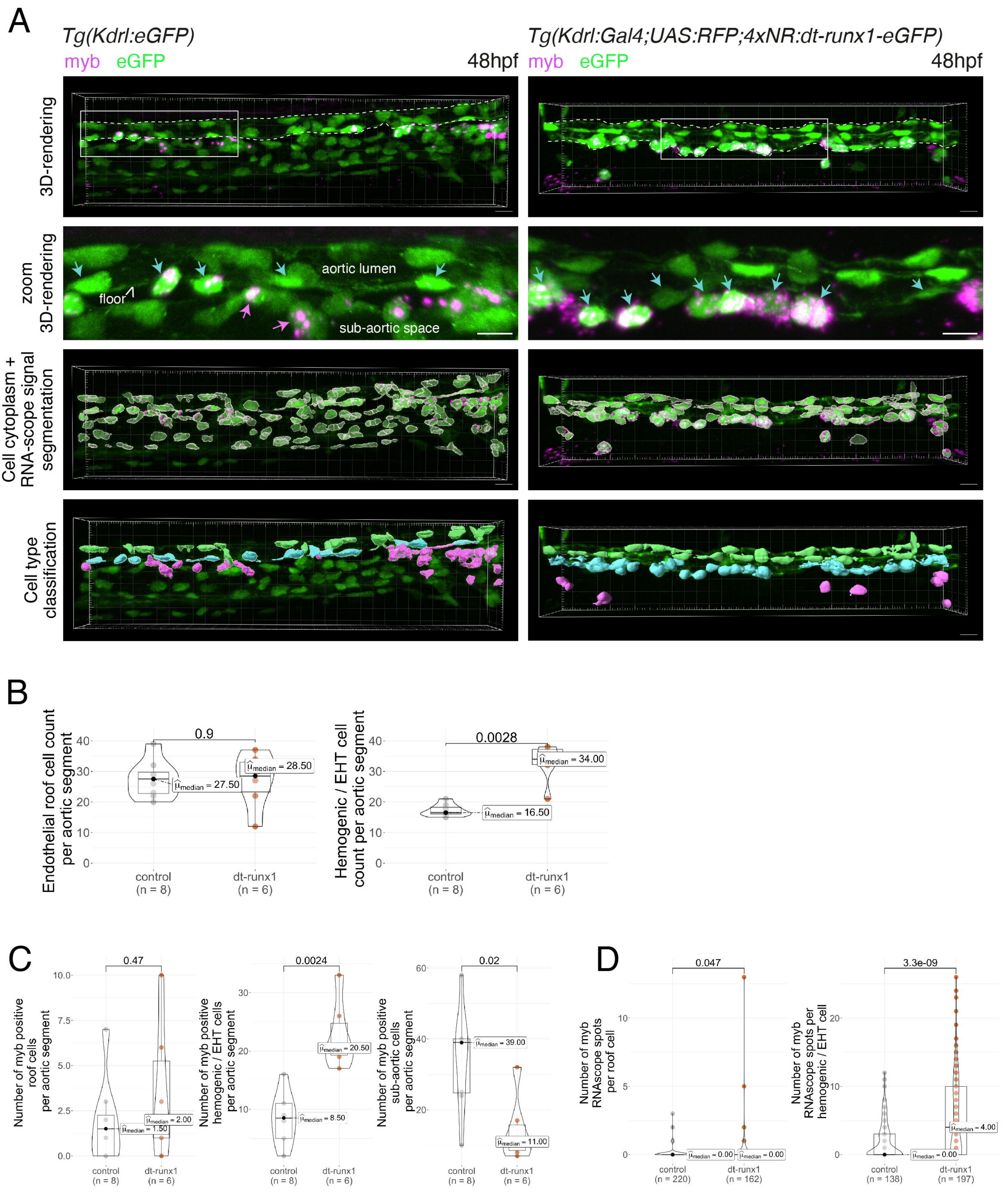

### Figure 3 - figure supplement 2

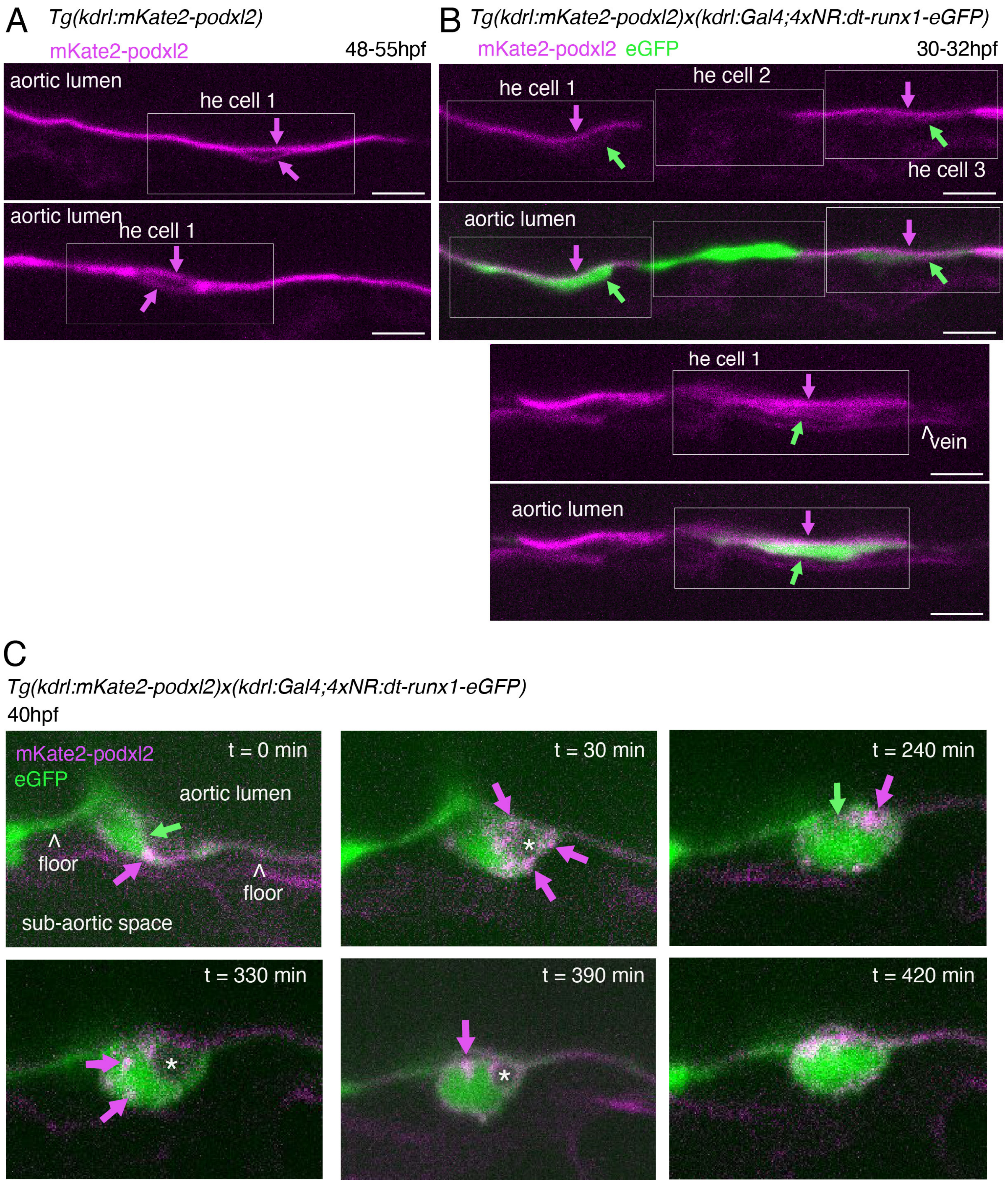

### Figure 3 - figure supplement 3

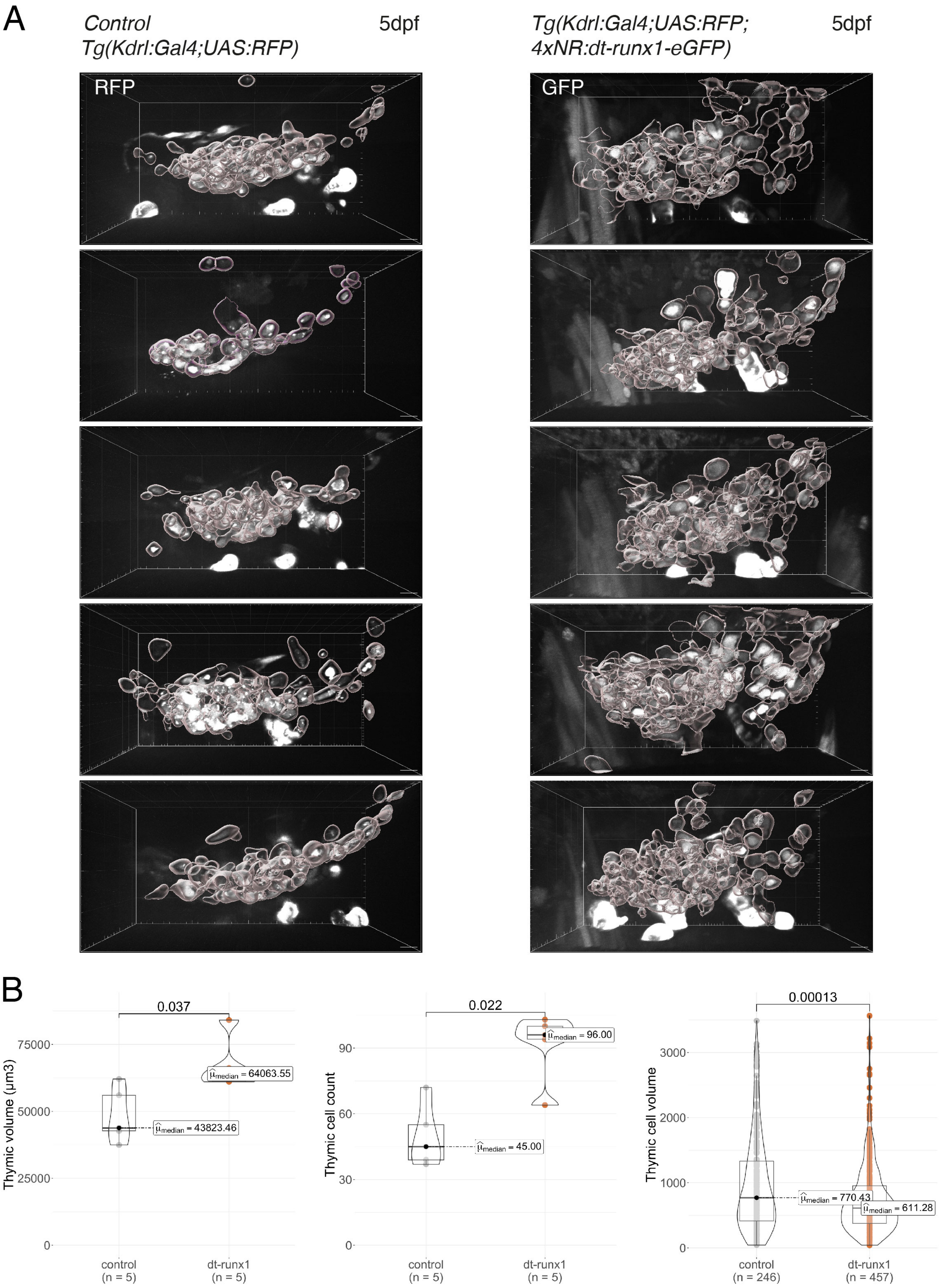

### Figure 4 - figure supplement 1

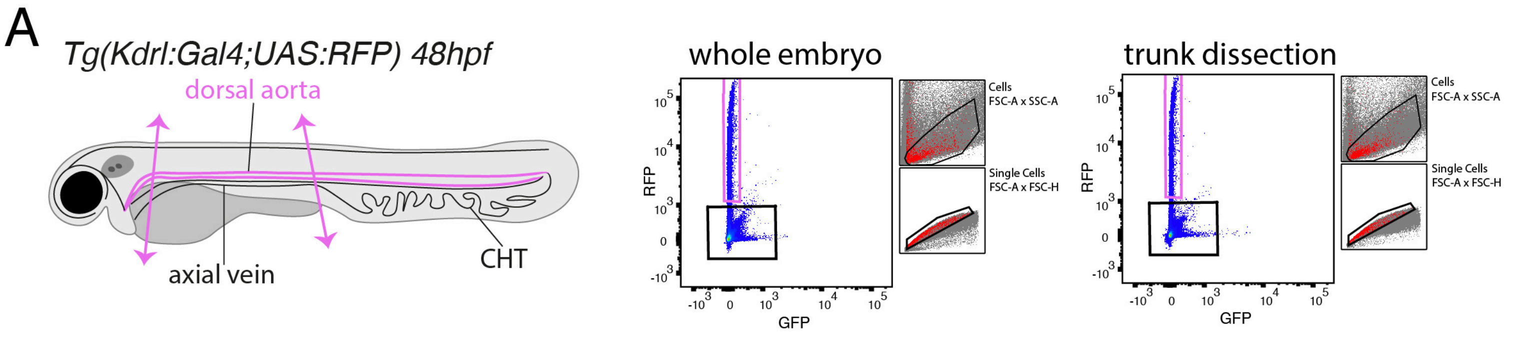


| | Whole embryo | | Trunk | |
|---|---|---|---|---|
| **RFP-** | 99.0 ± 0.0 | (n=2) | 97.7 ± 0.6 | (n=3) |
| **RFP+** | 0.95 ± 0.01 | (n=2) | 2.10 ± 0.59 | (n=3) |
| | % | | % | |

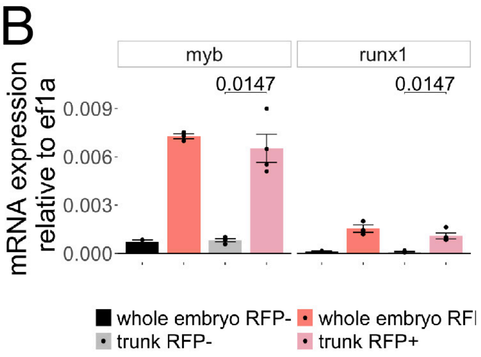


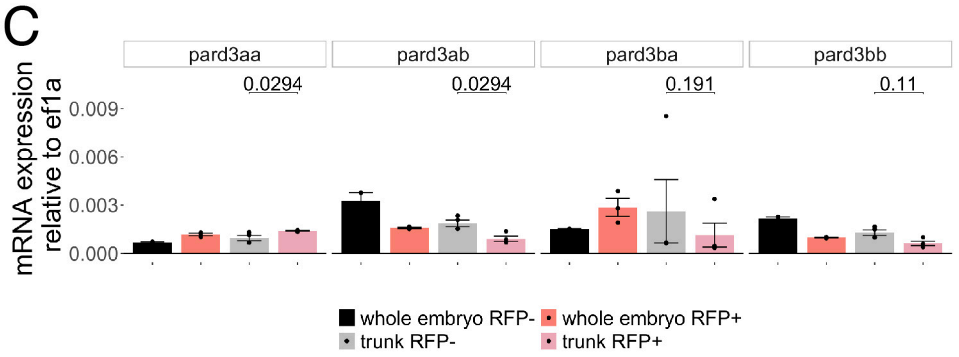

### Figure 4 - figure supplement 2

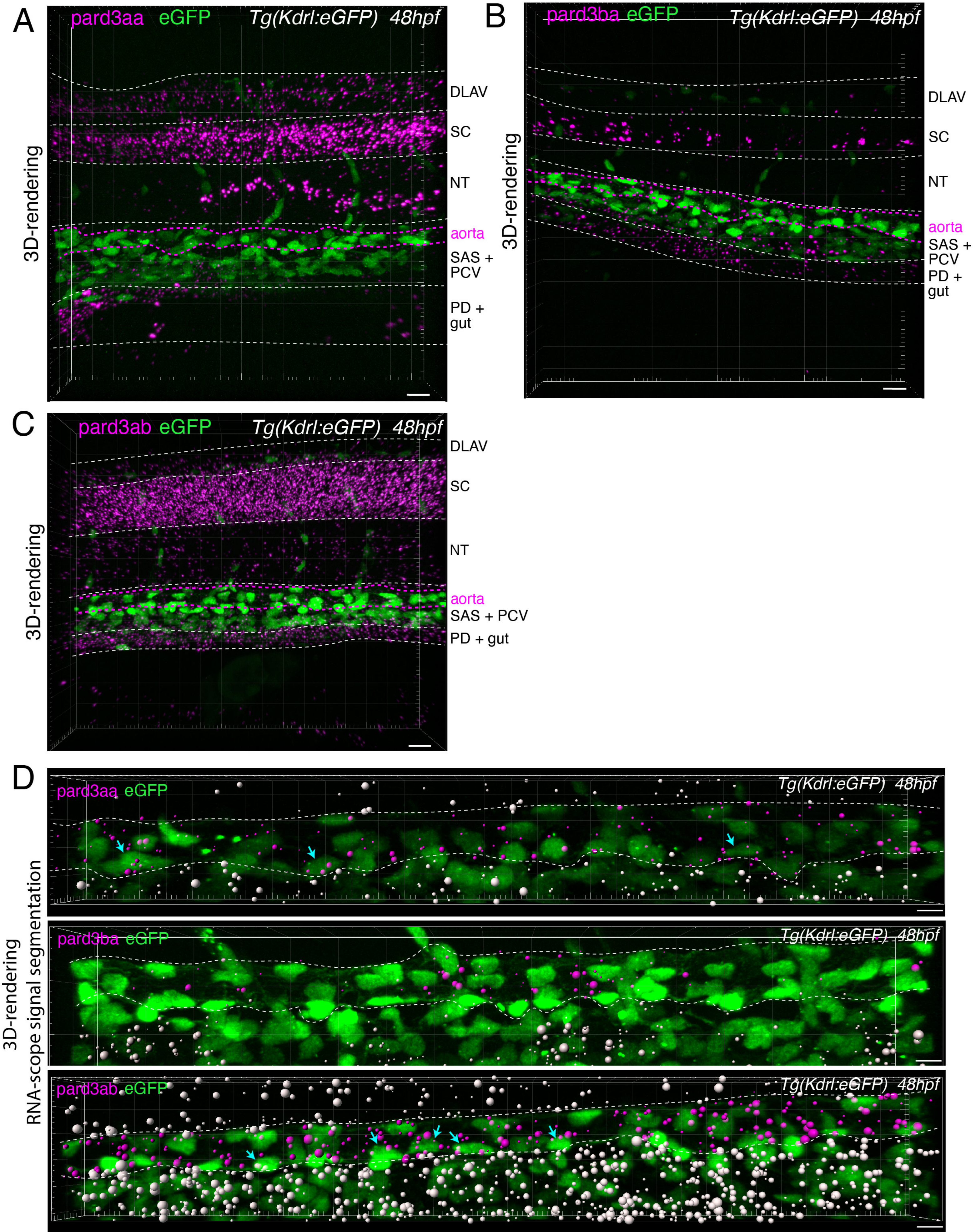

### Figure 4 - figure supplement 3

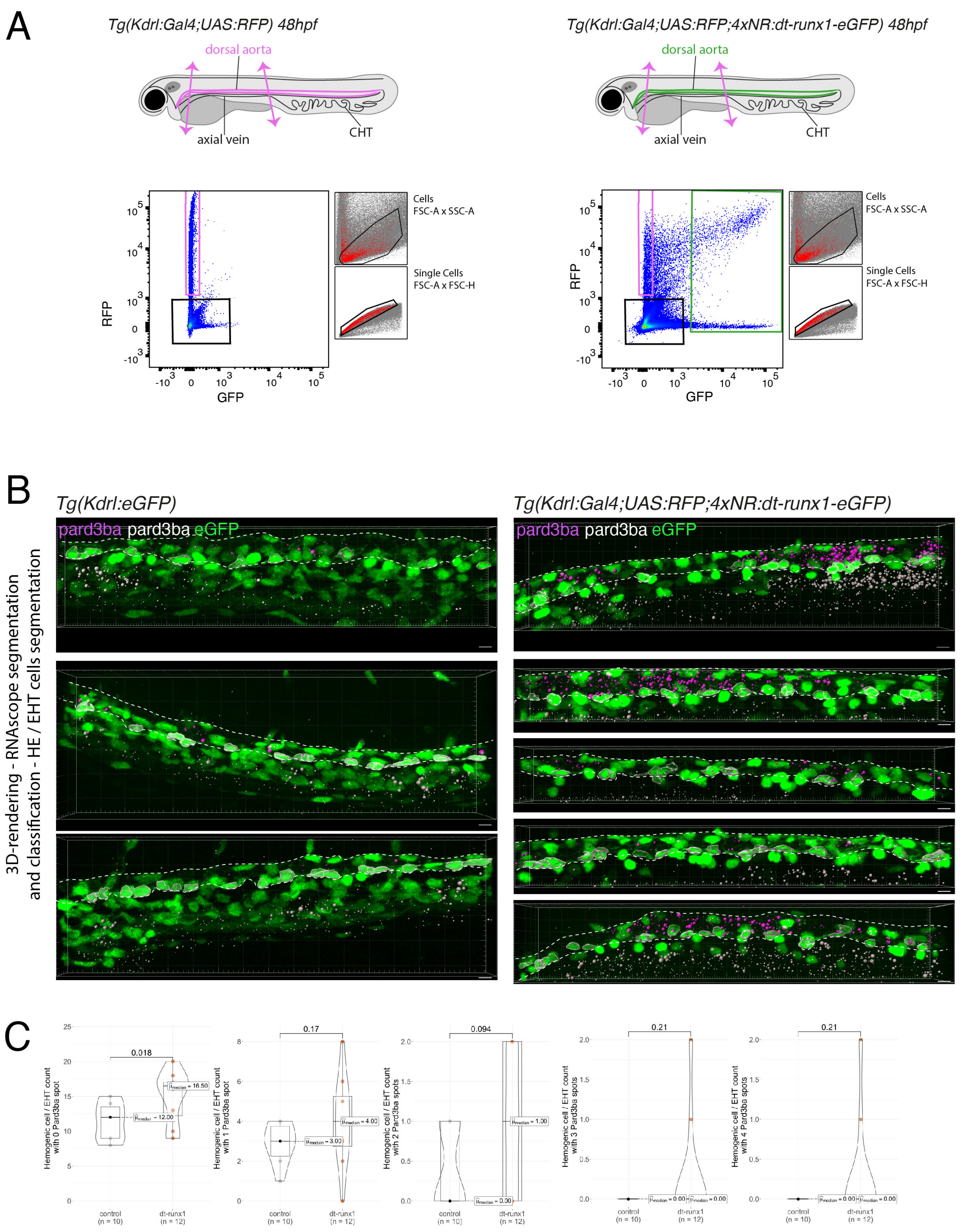

### Figure 4 - figure supplement 4

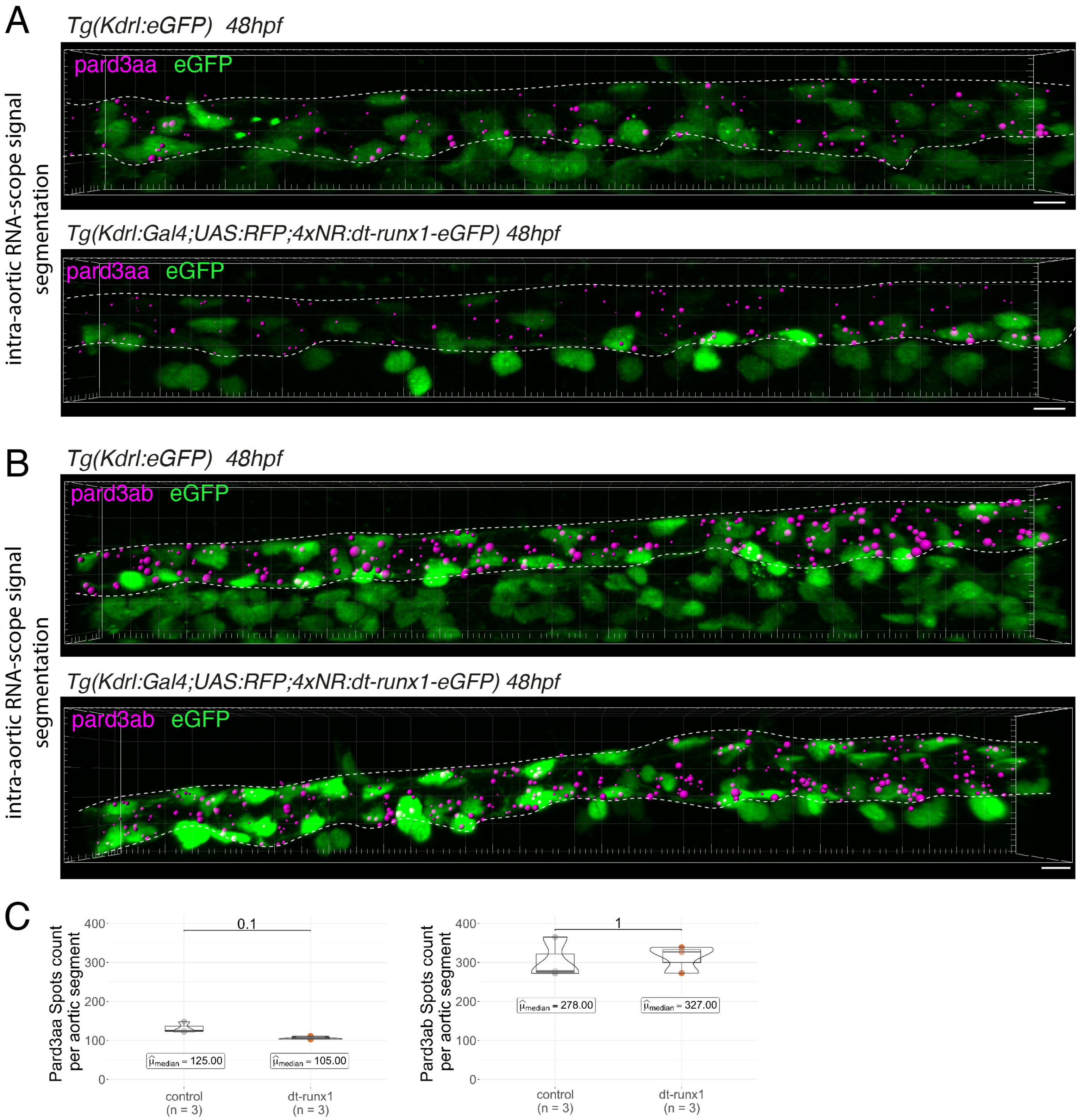

Figure 5 - figure supplement 1

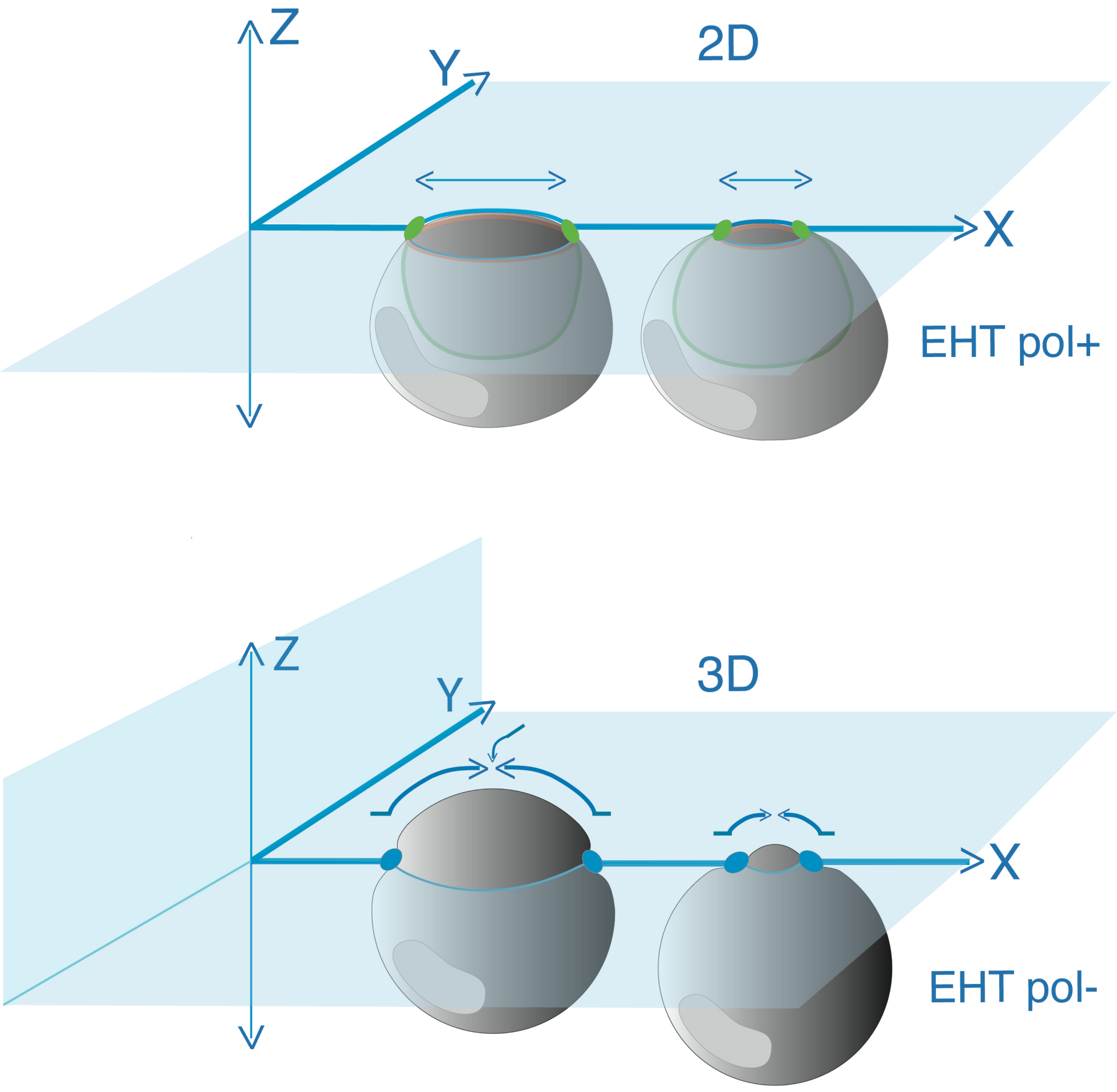

### Figure 5 - figure supplement 2

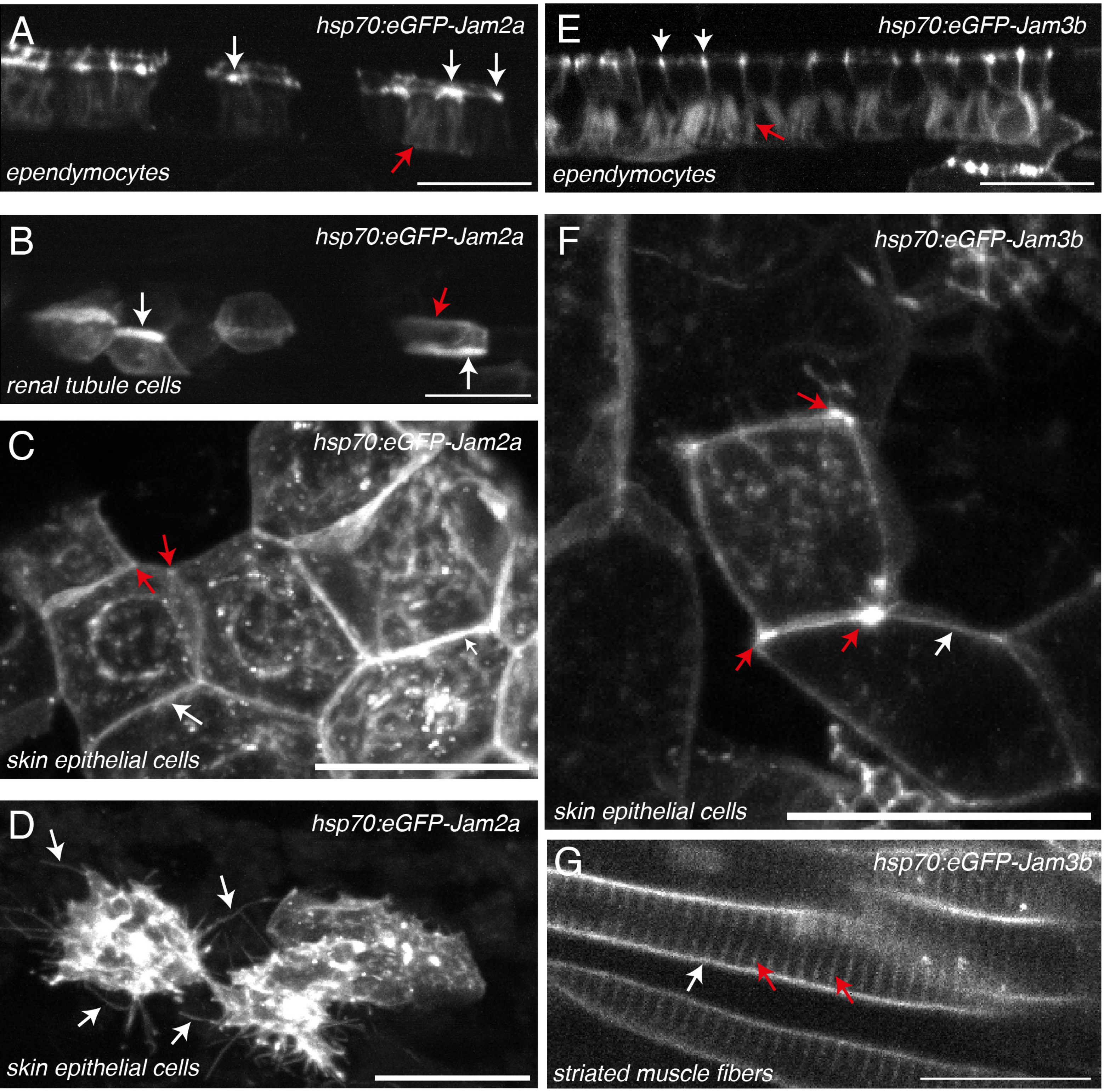

Figure 6 - figure supplement 1

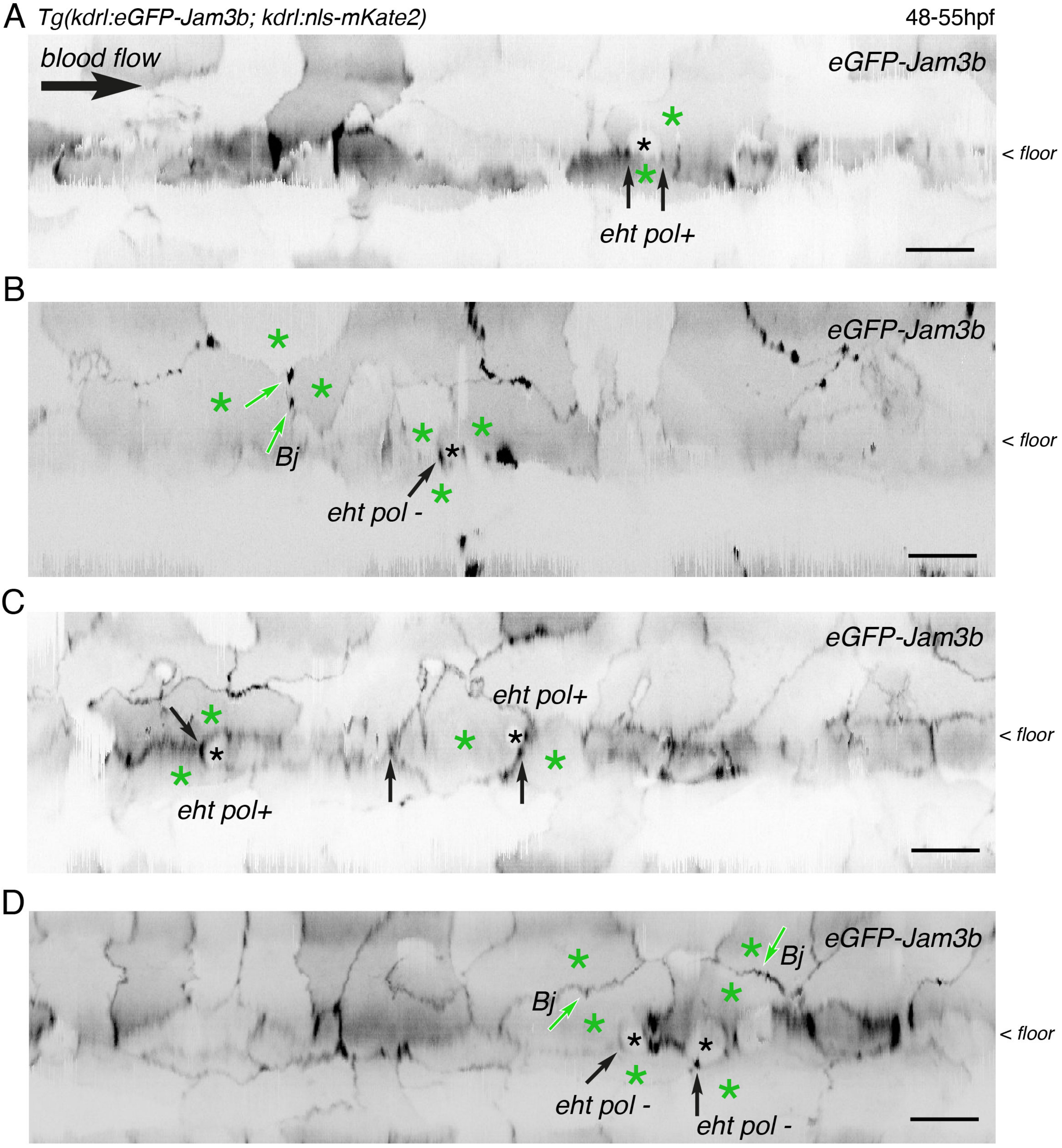

### Figure 7 - figure supplement 1

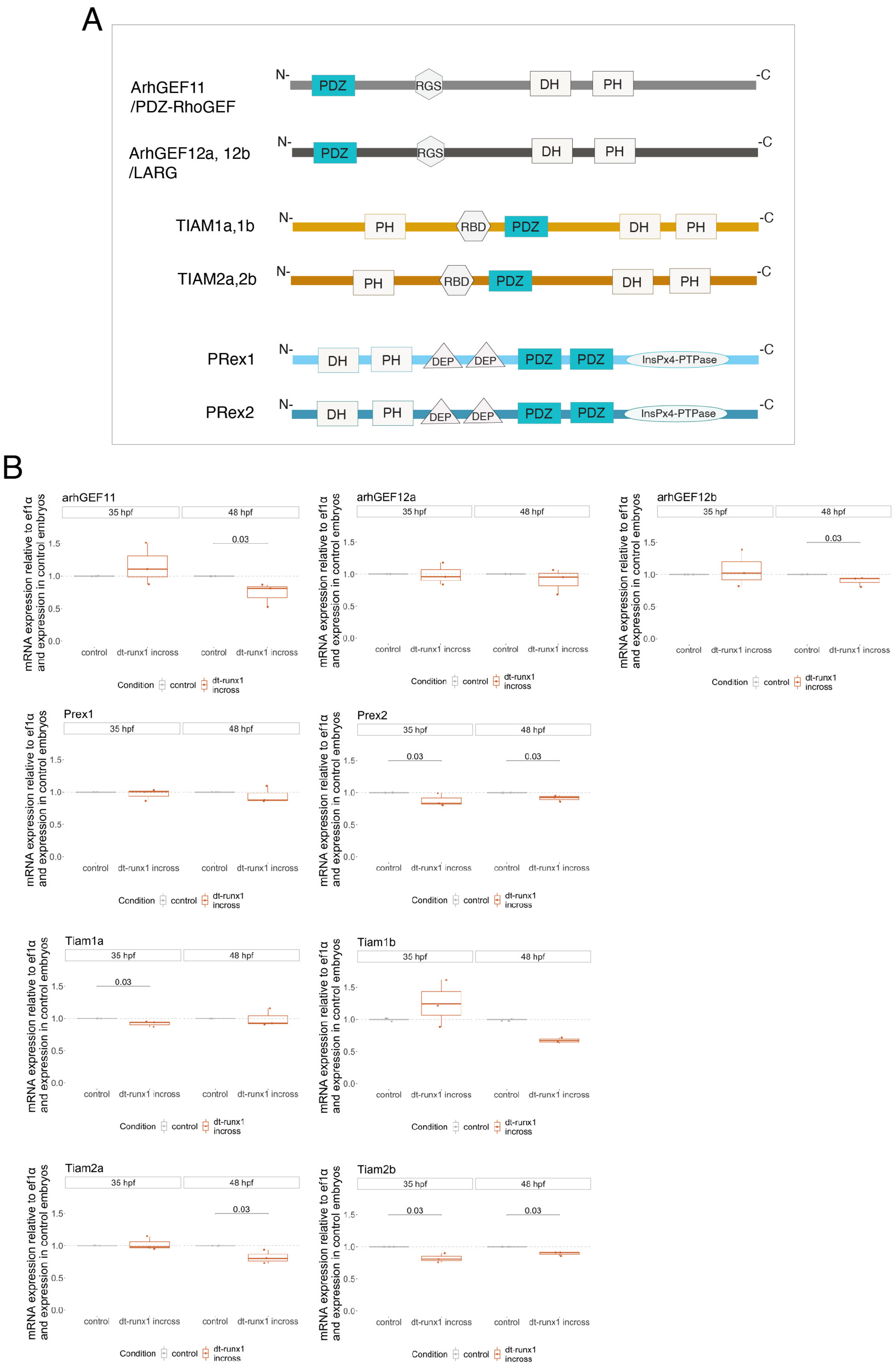

### Figure 7 - figure supplement 2

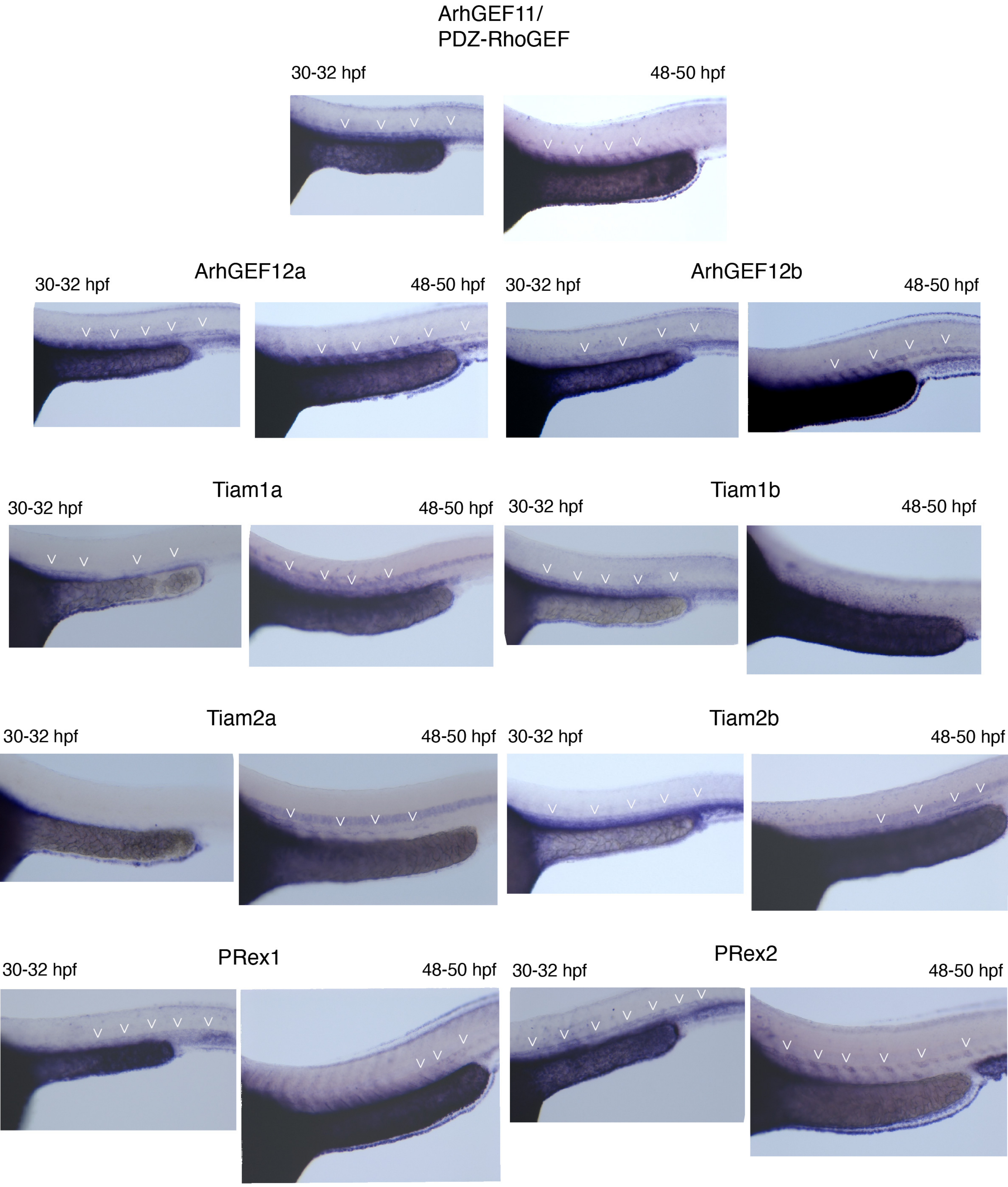

### Figure 7 - figure supplement 3

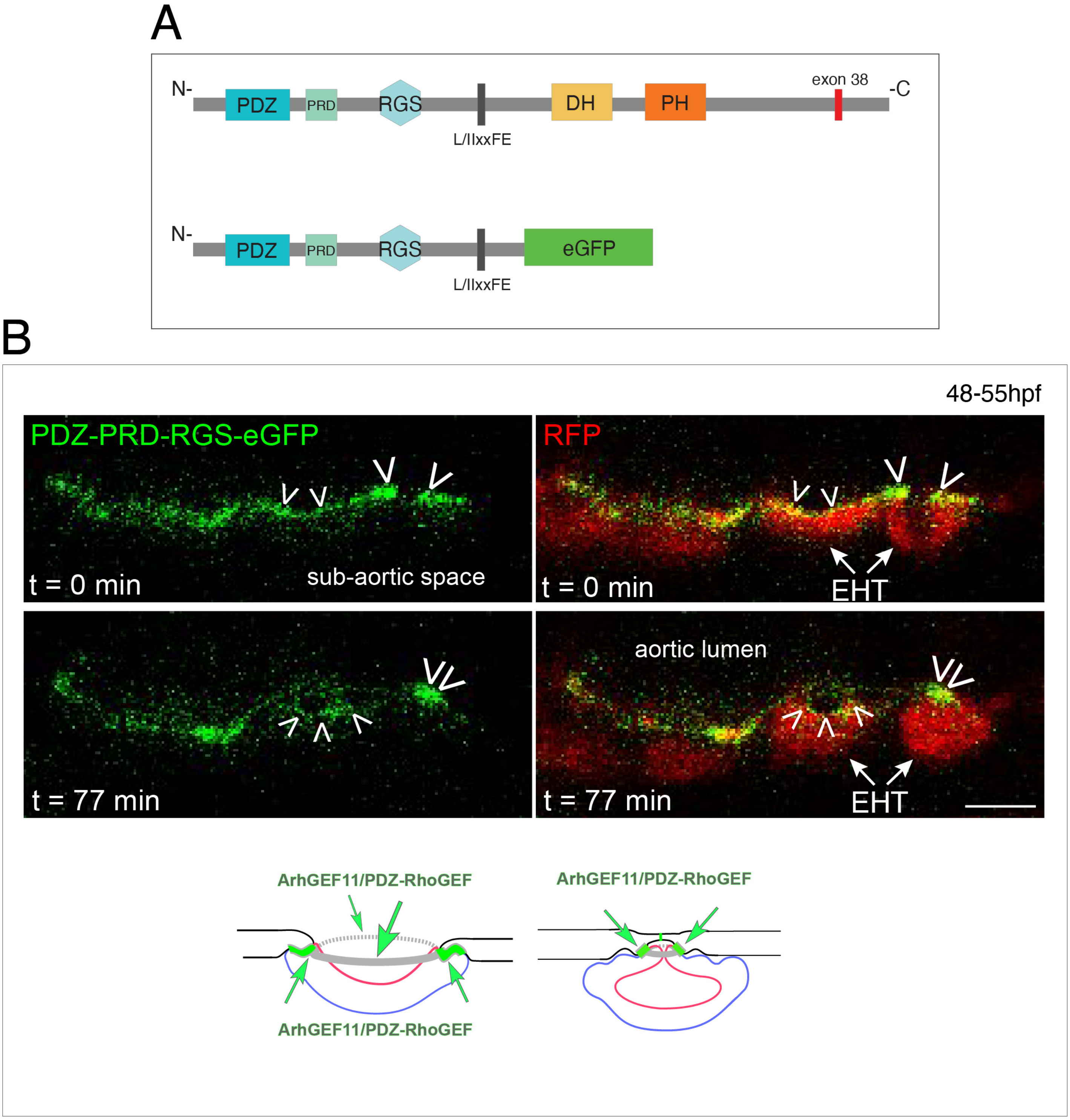

### Figure 7 - figure supplement 4

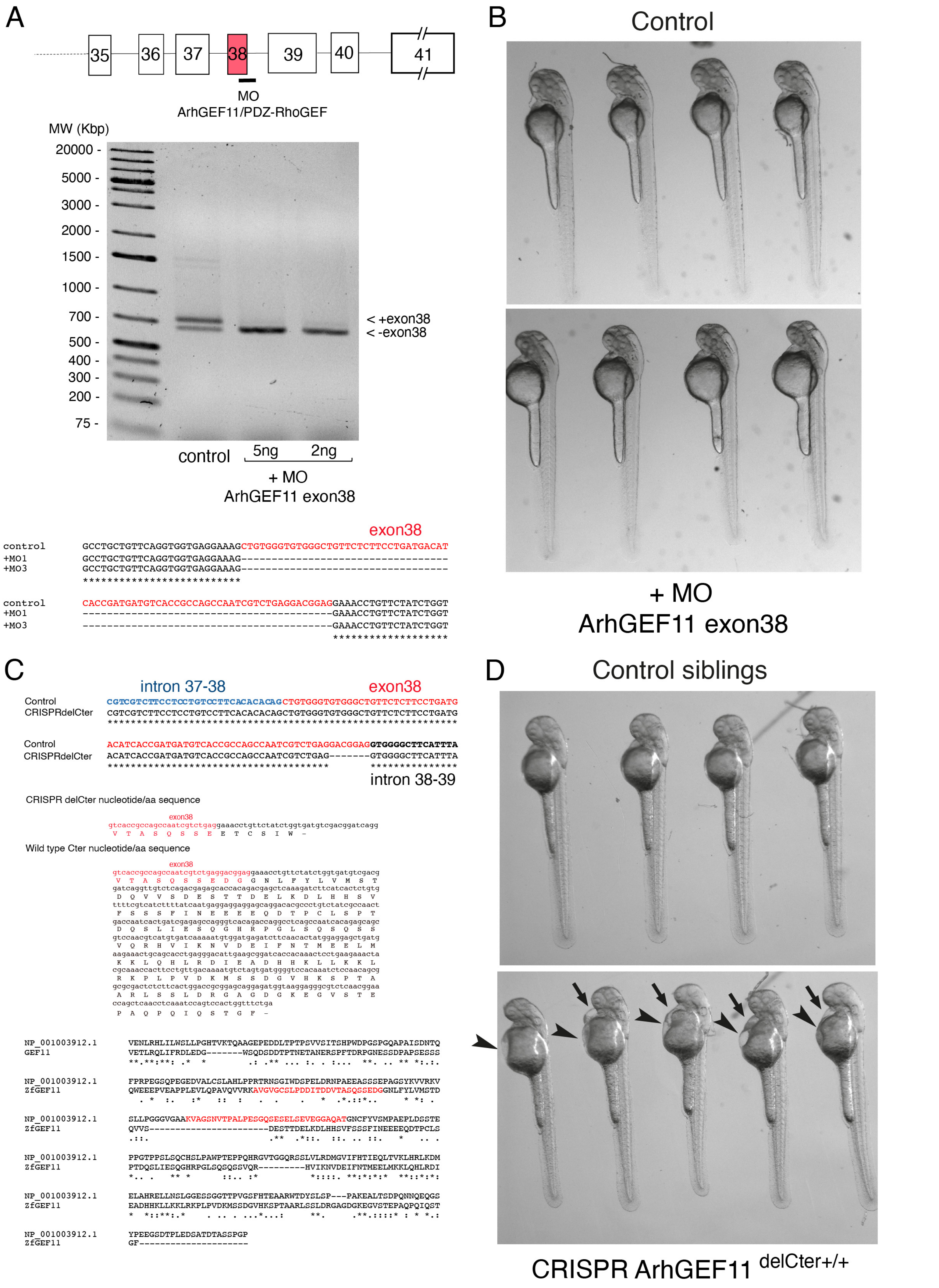

### Figure 7 - supplemental 5

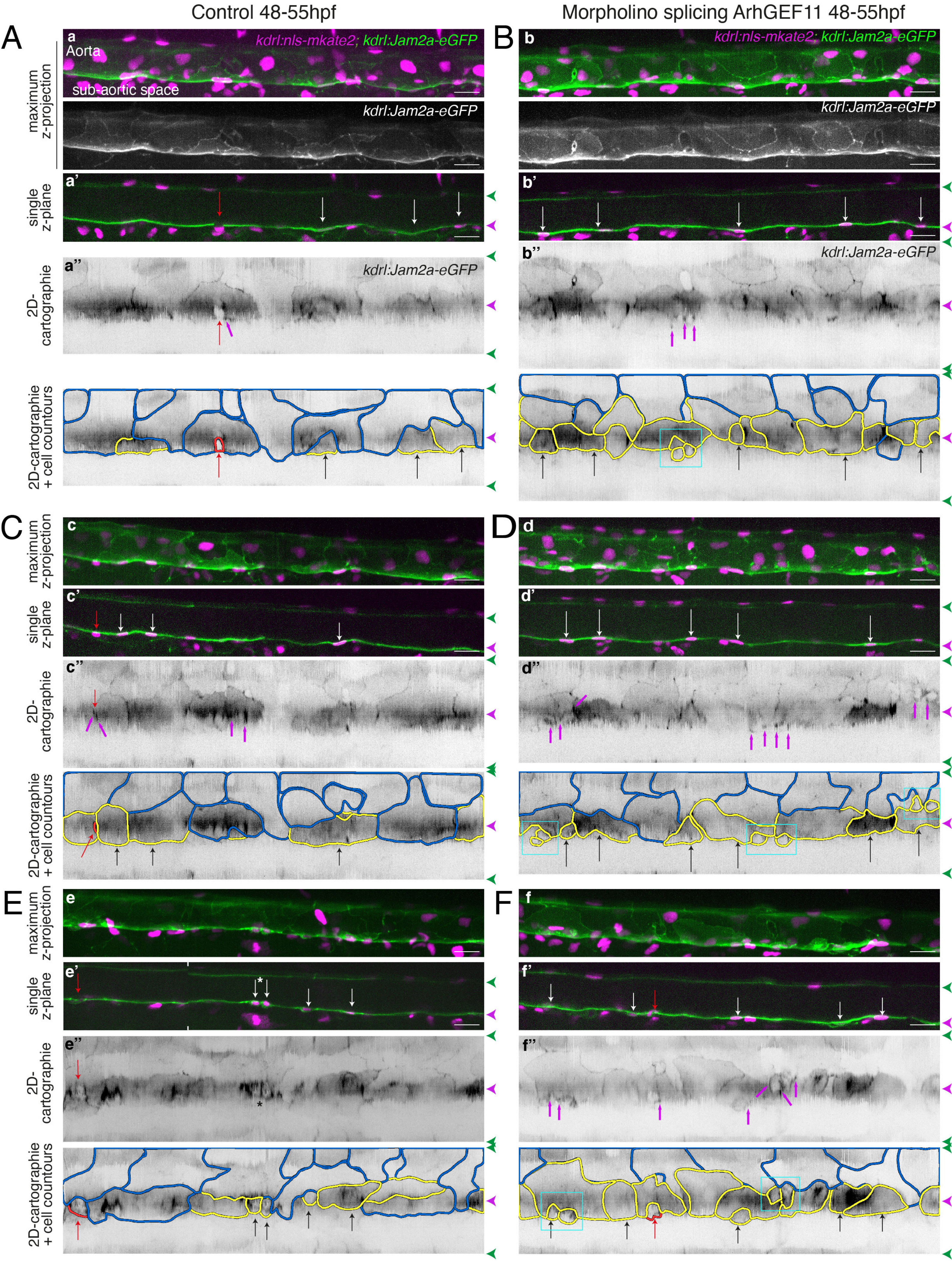

### Figure 7 - supplement 6

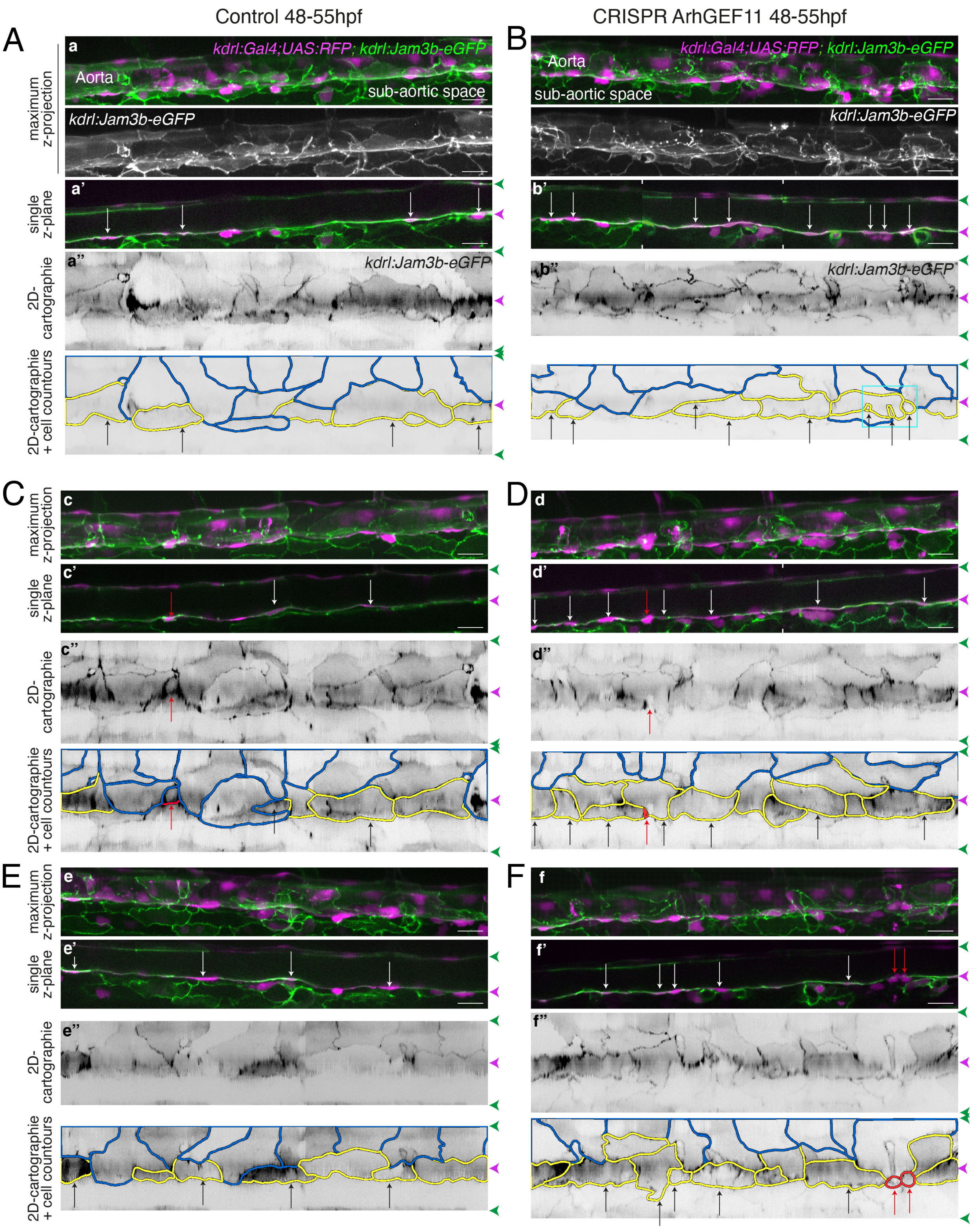

### Figure 7 - figure supplement 7

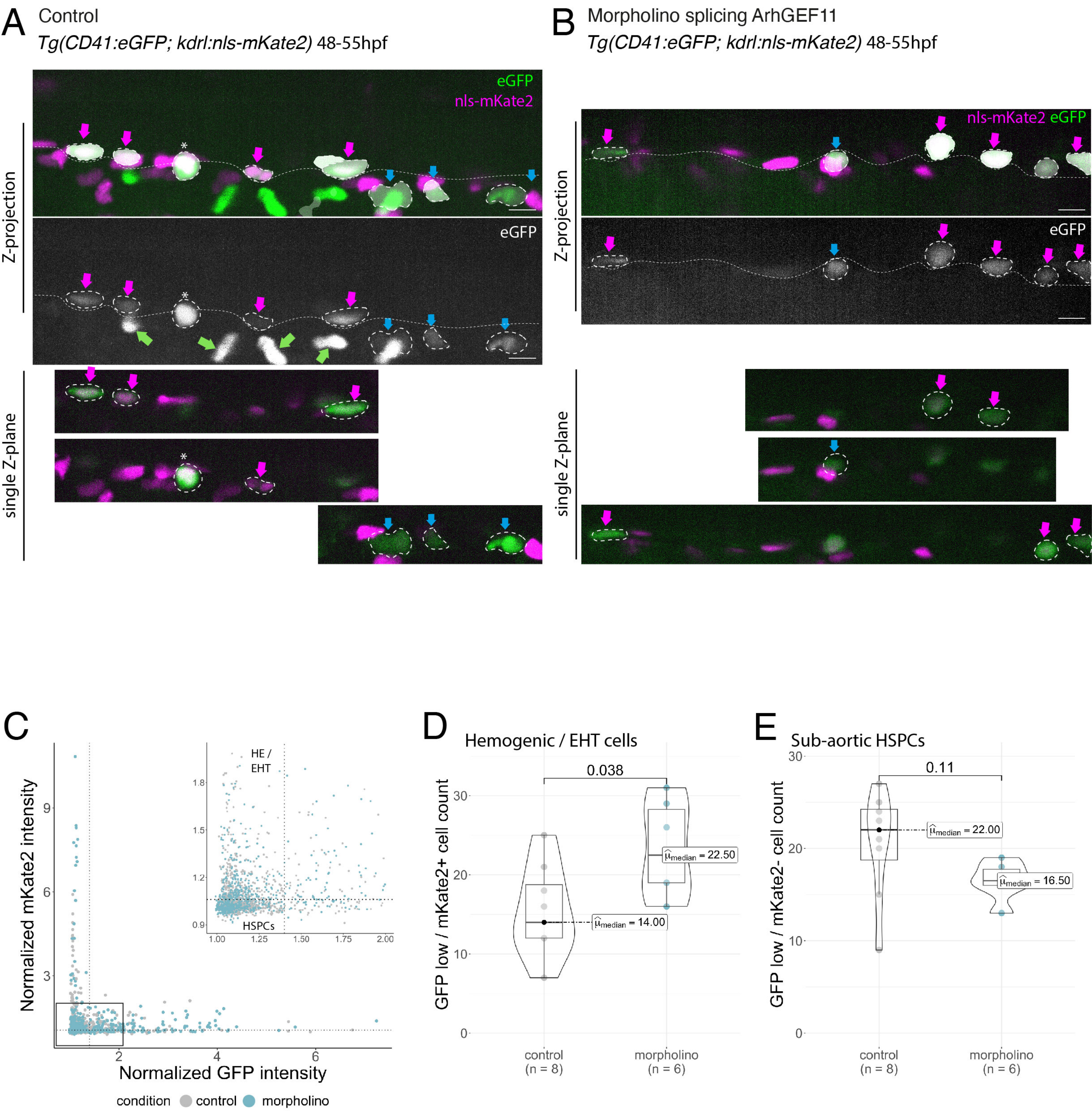
